# Plant genetic and root-associated microbial diversity modulate *Lactuca sativa* responsiveness to a soil inoculum under phosphate deficiency

**DOI:** 10.1101/2025.04.02.646582

**Authors:** Arianna Capparotto, Alessandro Ciampanelli, Paolo Salvucci, Guillaume Chesneau, Johannes Herpell, Simone Sello, Cristina Sudiro, Pieter Clauw, Adriano Altissimo, Stéphane Hacquard, Francesco Vuolo, Marco Giovannetti

## Abstract

- Microbial-based approaches have been proposed as a solution to decrease the use of chemical fertilizers in agriculture. Among these, the most promising candidates are arbuscular mycorrhizal fungi (AMF), with their ability to extend the root surface and absorb phosphate, and phosphate solubilizing bacteria (PSB), but their effectiveness has been shown to depend on plant genetic diversity.
- With the aim of identifying genetic markers explaining plant differential responses to soil-beneficial microbes, we monitored a panel of 128 fully sequenced varieties of *Lactuca sativa* under controlled P starvation conditions, treated with AMF and PSB.
- Results showed a strong effect of the lettuce genetic variation on the plant physiological and morphological response to the inoculum. Through genome-wide association studies, we identified specific genetic regions associated with variations in leaf phosphate and shoot biomass in response to the treatment.
- Beyond genetic factors, we detected changes in fungal β-diversity and increases in bacterial α-diversity associated with phenotypic variation, and we identified 44 ASVs linked to variation in agriculturally important traits. Among these, we experimentally validated the role of six bacterial strains through both in vitro and pot experiments, in affecting the leaf phosphate concentration and plant shoot biomass.
- In conclusion, we highlighted key genetic and physiological mechanisms that could play a crucial role in enhancing microbial treatments for optimizing plant phosphate management.

## 1. Introduction

Phosphate is one of the essential macronutrients for plants, as it plays a role in many enzymatic reactions and metabolic pathways and is a key component of macromolecules like nucleic acids, ATP, and phospholipids (Chiu & Paszkowski, 2019). Although soil contains approximately 0.05% (w/w) of phosphate, only about 0.1% of this is readily available to plants due to its limited diffusion rate and tendency to be bound in inaccessible organic forms (Zhu *et al*., 2011; Sharma *et al*., 2013; Iftikhar *et al*., 2024; Kaur *et al*., 2024). As a result, phosphate is the second most common limiting macronutrient for plant growth, after nitrogen.

To address this limitation, modern agriculture still heavily depends on chemical fertilizers derived from mined rock phosphate reserves (Cordell *et al*., 2009). However, while synthetic phosphate fertilizers have helped increase crop yields, their overuse has had a negative impact on both plants and the environment. These effects include soil eutrophication, a decline in soil biodiversity, and a reduced ability of certain plants to establish symbiotic interactions, such as those with arbuscular mycorrhizal fungi (AMF) (Martín-Robles *et al*., 2018) and nitrogen-fixing bacteria (Kiers *et al*., 2007; Liu *et al*., 2020). Additionally, the gradual depletion of finite phosphate reserves has driven up fertilizer costs. Together, these issues highlight an urgent need for sustainable agricultural strategies that reduce reliance on chemical fertilizers.

An eco-friendly approach gaining attention since the start of this century shifts the focus from nutrient inputs to the role of microorganisms (Capparotto & Giovannetti, 2023; Giovannetti *et al*., 2023). Throughout evolution, plants have developed beneficial associations with bacteria and fungi, particularly AMF and phosphate-solubilizing bacteria (PSB), as their primary strategy to cope with phosphate starvation in soils (Javot *et al*., 2007; Kaur *et al*., 2024). These associations are essential for plant health and provide key ecosystem services that sustain soil health. Nevertheless, modern breeding has often prioritized traits like biomass and yield, reducing plants’ responsiveness to forming symbiotic relationships (Alaux *et al*., 2024).

Thus, developing synthetic microbial communities capable of persisting in soil and functioning effectively in open-field conditions, alongside cultivating plant genotypes that respond efficiently to interactions with these microorganisms, remains a critical focus in research and agriculture (dos Reis *et al*., 2024). Indeed, while the effectiveness of nitrogen-fixing bacteria is well-consolidated and widely applied in agriculture (Liu-Xu *et al*., 2024; Thakur *et al*., 2024), much research is still needed to develop effective microbial inoculants that facilitate phosphate assimilation, especially in open-field conditions (O’Callaghan *et al*., 2022). For PSB, although various studies have shown a positive impact on nutrient uptake and crop yield, this effect is often achieved only when PSB are supplemented alongside reduced doses of chemical fertilizers (Suleman *et al*., 2018; Javeed *et al*., 2019).

Additionally, there are concerns about whether they can be used across a broad range of plants or if their effectiveness is limited by species-specific or genotype-specific affinities. Indeed, it is well-established that plant genotypes can influence responses to phosphate levels (Giovannetti *et al*., 2019) and bacterial recruitment in various plant compartments (Bulgarelli *et al*., 2013a; Capparotto *et al*., 2026). Moreover, numerous studies have highlighted the impact of intraspecific plant variation on associations with plant growth-promoting bacteria (PGPB), both in the model plant *Arabidopsis thaliana* (Plucani do Amaral *et al*., 2023) and in crops such as maize (Cuna *et al*., 2016; Vidotti *et al*., 2019; Meier *et al*., 2022) and rice (Edwards *et al*., 2015; Maghboli Balasjin *et al*., 2022). However, a clear understanding of the underlying genetic factors driving variation in plant-beneficial microbe interactions remains elusive.

In addition to the plant’s ability to assimilate phosphate, the integration of other morphological parameters is essential for gaining a comprehensive understanding of the plant response to microbial inoculation. Therefore, understanding whether different phenotypic responses are linked to distinct bacterial communities in the roots is crucial for identifying advantageous combinations of genetic regions, microbial consortia, and phenotypic traits that contribute to the environmentally sustainable development of resilient plants in agriculture (Escudero-Martinez & Bulgarelli, 2023).

In this study, we explored the role of intraspecific variation among 128 *Lactuca sativa* genotypes in determining their responses to a soil microbial inoculum. To gain a comprehensive understanding of the plant’s reaction to this treatment, we integrated 15 plant morphological traits. Operating under the hypothesis that plant genotype influences the outcome of plant-microbial associations, we identified putative genomic loci associated with the observed diversity in plant responses to the microbial inoculum. Additionally, we observed that plant responses to the inoculum were associated with changes in root fungal and bacterial diversity, and identified bacterial amplicon sequence variants (ASVs), likely originating from the inoculum, linked to these beneficial effects. Among the identified ASVs, we isolated six strains and validated their causal effects on variation in leaf phosphate concentration and plant shoot biomass under two different experimental conditions. Apart from the direct effects of each of these communities, we observed a potential correlation between the fungal phylum Glomeromycota and five bacterial ASVs involved in regulating key nutritional parameters. This suggests that fungi, through their intraradical and extraradical hyphal network, may represent a niche for bacteria with significant effects on plants.

Our findings provide a foundation for understanding the genetic and phenotypic basis of lettuce-soil microbe interactions, particularly in facilitating nutrient acquisition, and guide the identification and isolation of new effective microbes.

## 2. Material and methods

### 2.1. Experimental design

In this study, 131 fully-sequenced *Lactuca sativa* (lettuce) genotypes were provided by the Centre for Genetic Resources (CGN, Wageningen, Netherlands, Wei *et al*., 2021). Plant germination and growth took place in greenhouse conditions at LandLab S.r.l. in Quinto Vicentino, Vicenza (Italy). Before starting the main experiment, a pilot study was conducted to determine the optimal phosphate concentration that would induce a state of starvation without compromising the plants’ viability (Supplementary Table **1**). 131 lettuce genotypes were sown in seed trays in 12 replicates each and germinated in a growth chamber for approximately one month. During this initial growth phase, the plants were fertigated with a nutritional solution containing NPK 18/18/18 to support their development. To simulate phosphate starvation, plants were grown in sand as a medium and supplemented with triple superphosphate (TSP) at a rate of 32.61 kg/ha (or 0.33 g per plant), equivalent to 15 kg/ha of P₂O₅.

The procedure involved filling each pot (15 cm x 15 cm x 20 cm) with 3 L of non-sterilized soil-sand mixture, adding the pre-weighed TSP, and then topping it up with an additional 1 L of soil-sand to complete the pot-filling process. A mesh pad (30 g/m²) with holes was placed at the base of each pot to prevent water spillage. Five weeks after germination, a total of 866 young seedlings were transplanted into the previously prepared pots in the greenhouse and randomized positions as outlined in Supplementary Table **2**. Three genotypes (TKI-93, TKI-23, TKI-121) that did not germinate sufficiently to achieve the required number of replicates for the experiments were excluded from the analysis, resulting in a total of 128 genotypes being considered. Four replicates per genotype were treated with 50 mg of microbial inoculum, applied as a 50 ml aqueous solution directly to the plant rootlet, at a concentration comparable to that used under field conditions. The synthetic microbial community used was formulated by Sacco System, located in Cadorago, Como (Italy). This community comprised two strains of arbuscular mycorrhizal fungi (*Rhizophagus irregularis* and *Funneliformis mosseae*) in a concentration of 1000 propagules/g, and two strains of phosphate-solubilizing bacteria (*Bacillus subtilis* and *Chryseobacterium idoltheticum*) in a concentration of 1 × 10^9^ CFU/g each. These microorganisms were isolated for their capacity to enhance phosphorous availability and promote plant growth. Because arbuscular mycorrhizal (AM) fungi are obligate symbionts, their propagules were introduced in the form of root fragments enriched with the desired fungal species, which may also contain additional fungal and bacterial taxa. The microbial inoculum had a shelf life of two years, as stated by the company, thereby covering the entire duration of the experiment (three months). To support plant growth, periodic fertigation was implemented at a concentration of N-K_2_O 250, equivalent to 76.92 kg/h. Detailed information regarding dates and concentrations of fertigation are included in Supplementary Table **3**.

### 2.2. Plant phenotyping

Throughout the entire experiment, plant growth, phenotypic appearance, and physiology of each plant were recorded twice daily using a non-invasive, high-throughput phenotyping platform, the PlantEye (PlantEye F600, Phenospex, Heerlen, The Netherlands), provided by the LandLab company. The analysis combined 3D imaging with multispectral imaging. In this study, the following parameters were analyzed (Zieschank & Junker, 2023):

1. Greenness/Green Leaf Index (no units, Equation 1): The ratio of green reflectance to red and blue reflectance. It indicates the combined effects of leaf physiology and canopy structure.

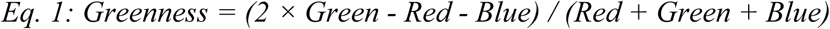
2. Hue (°): Represents the visible color determined by the wavelength of reflected light, independent of saturation and brightness. It indicates photosynthetic pigment levels and nutrient concentration in leaves.
3. Digital Biomass (cm³, Equation 2): The volume of the plant’s above-ground biomass, indicating plant productivity.

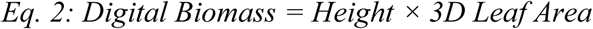
4. Height (mm): The average height of the community’s plants, from the ground to the top 10% of plants. It is an indicator of plant productivity, nutrient availability, light competition, and growth strategy.
5. Leaf Area (cm²): The three-dimensional leaf area, corrected for leaf inclination, indicating heat load, water retention, and gas exchange.
6. Leaf Area Index (cm²/cm², Equation 3): The leaf area per unit ground area, related to photosynthetic activity and respiration.

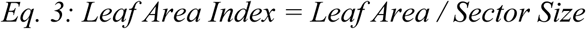
7. Light Penetration Depth (mm): The deepest point the laser can penetrate through the canopy, indicating vegetation density and resource availability.
8. Normalized Digital Vegetation Index (NDVI, no units, Equation 4): The ratio of near-infrared (NIR) to red reflectance, indicating vegetation type, productivity, and growth parameters like leaf area index and biomass.

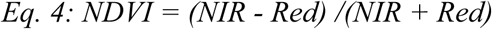
9. Normalized Pigments Chlorophyll Ratio Index (NPCI, no units, Equation 5): The ratio between the blue and red channels, associated with chlorophyll content and indicating the plant’s physiological state.

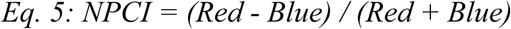
10. Plant Senescence Reflectance Index (PSRI, no units, Equation 6): The ratio of red and green to near-infrared reflectance, associated with carotenoid and chlorophyll content, indicating leaf senescence.

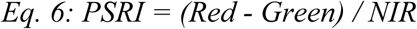

Since daytime measurements could be influenced by various confounding factors only the nighttime recordings were used for the analysis presented in this paper. The scanning sequence for PlantEye is detailed in Supplementary Table **4**, and measurements for each plant replicate are provided in Supplementary Table **5**.

### 2.3. Plant sampling

Leaf sample collection took place 5 weeks after transplanting (December 14 and 15, 2022). The procedure involved weighing the aerial part of the plant before leaf sampling with a portable balance (sensitivity=0.01 g). Each leaf was sampled using a single-hole punch to obtain three leaf disks with a diameter of 3 mm from each leaf. These leaf disks were then combined with those obtained from two additional leaves of the same plant, resulting in a total of 9 leaf disks per replicate. Simultaneously, roots were washed to eliminate soil residues, dried using absorbent paper, and weighed with the same portable balance to obtain their fresh weight. The root and shoot fresh weights are reported in Supplementary Table **6**. Subsequently, all treated roots and a few control ones were sampled for metabarcoding analysis. Specifically, a bundle of approximately 2 cm in length roots (excluding the primary roots) was collected. Both leaf and root samples were gathered in racked 96 collection microtubes (QIAGEN Hilden, Germany), following the order outlined in Supplementary Table **7**. In addition to these, the soil used for plant germination and the sand used in the experiment mixed with the water used for watering the plants were also collected and used as controls. The tools used for sampling were washed with 70% ethanol and distilled water between each accession. All the collected materials, except for the soils, were stored at −80°C before undergoing lyophilization for long-term storage.

### 2.4. Inorganic phosphate quantification

Lyophilized leaf disks were weighed using an ultra-analytical balance (sensitivity = 0.01 mg) to achieve a final sample weight between 0.25 and 0.80 mg. The disks were then freeze-dried and ground into a fine powder using a TissueLyzer (Qiagen, Hilden, Germany) before being resuspended in NanoPure water and incubated at 98°C for 1 hour. The final suspension, obtained after centrifugation, was used for sample processing with the phosphate assay kit (MAK308, Sigma-Aldrich, St. Louis, USA), following the instruction manual with some modifications. Briefly, 16 µL of the sample suspension (and separately 16 µL of the standard solution) was mixed in replicates with 32 µL of Malachite Green Reagent. After a 30-minute incubation period, absorbance was measured at 620 nm using a Tecan spectrophotometer (Trading AG, Männedorf, Switzerland). Using the absorbance and intercept values, we calculated the inorganic soluble phosphate concentration, expressed as µmol/g. The phosphate quantification table is detailed in Supplementary Table **8**.

### 2.5. Plant phenotypic response and metabolites quantification

For each parameter measured by Phenospex, the overall plant response to the inoculum was determined by calculating the ratio between the median values of the treated and control samples for each genotype. The data was then visualized in R by applying z-scaling and creating a heatmap using the pheatmap package (version 1.0.12), as well as through principal component analysis (PCA) with the FactoMineR (version 2.11) and factoextra (version 1.0.7) packages. From the representation of the first and second principal components, we identified four groups of genotypes that differently responded based on their positions at the extremes of the phenotypic space and their locations within the four quadrants around the analysis vectors. This process led to the selection of 28 genotypes showing contrasting responses, which were subsequently confirmed through metabolite quantification using the rainbow protocol, with minor modifications (López-Hidalgo *et al*., 2021). In brief, lyophilized leaf disks were weighed using an ultra-analytical balance (sensitivity = 0.01 mg) to ensure a final sample weight of at least 1.30 mg. The disks were then freeze-dried, ground into a fine powder using a TissueLyzer, and suspended in 600 µL of 80% ethanol. For photosynthetic pigments (PHP) quantification, 200 µL of the supernatant was diluted 1:2. All subsequent steps for free amino acids (FAA), total soluble sugars (TSS), and starch (STA) analyses were performed according to the published protocol. Their final concentrations were expressed in µL/mL. The metabolite quantification table is detailed in Supplementary Table **9**.

### 2.6. Genome-wide association mapping

To associate plant physiological variation in lettuce, we performed a Genome-Wide Association Study (GWAS) using the *Lactuca sativa* SNP matrix. Analyses were conducted in Visual Studio Code (version 1.94.2) with the GEMMA software package (version:0.98.5, Zhou and Stephens, 2012), employing a linear mixed model. Phenotypic traits were defined as the plant response to microbial treatment, calculated as the ratio between the median trait values of treated and control plants for each genotype. SNPs with a minor allele frequency (MAF) below 0.05 were excluded. To account for population structure, the first five principal components (PCs), obtained via PCA using PLINK2, were included as covariates in the model. A centered relatedness (kinship) matrix was computed in GEMMA using the standardized relatedness estimator and incorporated as a random effect to correct for genetic relatedness among genotypes. Association analysis was conducted with GEMMA’s linear mixed model applying restricted maximum likelihood (REML) estimation.

The full analysis script is available on Github (https://github.com/picla/salad_marco/tree/master/003.gwas/002.scripts). Broad-sense heritability for each trait was estimated in R using the ANOVA method.

### 2.7. DNA extraction and sequencing

Ninety-one root samples, including 6 controls and 85 treated samples from 22 genotypes, were selected for 16S and ITS sequencing based on their metabolite clustering and phenotypic responses to the inoculum (Supplementary Table **10**). In addition, four samples each from the inoculum, germination soil, and growth soil (sand + water) were included as environmental matrix controls. DNA extractions were performed using the CTAB protocol with some modifications. Briefly, approximately 3.5 mg of lyophilized root sample was ground using a TissueLyzer (Qiagen, Hilden, Germany) and then resuspended in 600 µl of CTAB, followed by incubation at 60°C for 1 hour. After this step, 600 µl of Phenol/Chloroform/Isoamyl alcohol (25:24:1) was added twice, mixing and removing impurities each time. The remaining supernatant was treated with ice-cold isopropanol and incubated at −20°C for 2 hours. DNA was then precipitated by centrifugation, washed with 70% ethanol, and resuspended in NanoPure water. Inoculum and sand with water samples were extracted using the same method, while the germination soil samples were extracted using the DNeasy® PowerSoil® kit (QIAGEN, Hilden, Germany) according to the manufacturer’s instructions. To further purify the samples and prepare them for PCR amplification, some were processed using the OneStep PCR Inhibitor Removal Kit (Zymo Research, Irvine, CA, USA). DNA from all 103 samples was quantified using Qubit™ dsDNA HS (High Sensitivity) Assay Kits from Thermo Fisher Scientific. Sample information and quantification values are detailed in Supplementary Table **11**. Subsequently, the DNA samples underwent amplicon sequencing of the 16S V5-V7 regions for bacteria and the ITS2 region for fungi using the MiSeq Illumina platform at NOVOGENE (UK). For bacteria, the V5-V7 region was amplified using the primer pair 799F (5′-AACMGGATTAGATACCCKG-3′) and 1193R (5′-ACGTCATCCCCACCTTCC-3′). For fungi, the ITS2 region was amplified with the primer pair ITS3F (5′-GCATCGATGAAGAACGCAGC-3′) and ITS4R (5′-TCCTCCGCTTATTGATATGC-3′).

### 2.8. Raw reads processing

Raw sequencing reads were processed using Dada2 (version 1.30.0) in R (version 4.3.3). For ITS analysis, only the forward reads were used as described by (Pauvert *et al*., 2019). Barcodes and primer sequences were removed before filtering, trimming, and merging the forward and reverse reads. The merged sequences then underwent chimera removal and taxonomic assignment using the Silva database (version 138, Callahan *et al*., 2016) for bacteria and the UNITE dynamic database (version 18.07.2023) for fungi. The dataset was further refined to remove chloroplast, mitochondrial, eukaryotic, and archaeal contaminants. For bacteria, only samples with a relative abundance greater than 0.01% were retained, reducing the number of ASVs from 45,316 to 35,043, while keeping all 103 samples. For fungi, samples with fewer than 100 counts and with a relative abundance larger than 0.01% were excluded, reducing the number of ASVs from 7,092 to 7,067 and removing five samples. After applying the same abundance threshold used for bacteria, the fungal ASVs were further reduced to 6,022. All bacterial samples underwent size-based rarefaction, set at 41,736 reads, matching the minimum read count among the samples. This process was performed in R with the use of vegan package (version: 2.6.4). For fungi, due to large differences between samples, rarefaction was performed per compartment: sand + water samples were rarefied at 42,893 reads, germination soil at 29,357, inoculum at 70,255, and root samples at 1000reads. This process reduced the number of root samples from 86 to 50, with a final total of 62 samples. The number of samples and ASVs lost at each step reported above is detailed in Supplementary Table **12**. Finally, to check for correlations between the fungal and bacterial communities, both root datasets were rarefied to the minimum sampling depth, corresponding to 1000 reads.

### 2.9. Evaluation of arbuscolar mycorrhizal colonization

The entire root systems of seven different genotypes (totaling 46 samples), chosen randomly during sampling, were collected and processed to assess the degree of arbuscular mycorrhizal colonization using the Trouvelot method (Trouvelot, 1986). Roots were first cleared by incubation in 10% KOH at 96 °C in a water bath for 10 minutes. Following incubation, the KOH solution was discarded, and roots were washed twice with 5% lactic acid. Subsequently, roots were stained in a solution containing 5% black ink and 5% lactic acid at 96 °C for 11 minutes. After staining, roots were washed repeatedly in 5% lactic acid until the solution became clear. From each biological replicate, five microscope slides were prepared, each containing 20 root fragments of approximately 1 cm in length. Therefore, around 100 cm of root tissue were analyzed per each sample.

### 2.10. Bacterial tests on lettuce plant growth

Six bacterial strains were selected from a representative collection of 105 lettuce root-associated isolates (doi:10.5281/zenodo.19664961) based on their genetic similarity to ASVs significantly associated with leaf phosphate content in lettuce roots (see Section 3.5). Briefly, the V4–V5 hypervariable regions of the 16S rRNA gene of the collection isolates were assembled into a single-fragment contig and aligned against the 22 responsive ASV sequences using VSEARCH v2.17.0 (Rognes et al., 2016), applying a minimum sequence identity threshold of 97%. Full alignment results, including percentage identity, query sequences, CIGAR notation, and taxonomic assignment, are reported in Supplementary Table **S13**. These strains were used to inoculate three lettuce genotypes (TKI-24, TKI-38, and TKI-100) in two experimental setups: a plate assay and a pot assay in a growth chamber.

Each strain was grown individually in 50% Tryptic Soy Broth (MP Biomedicals, cat: 1010717) for 48 h at 28°C, after which cultures were adjusted to a final OD₆₀₀ of 0.05. A synthetic community (SynCom) was generated by combining the six strains in equimolar proportions. As a mock control, a 10 mM MgCl₂·6H₂O solution (Merck, cat: 63033) was used.

For the plate assay, lettuce seeds were surface-sterilized by immersion in 0.5% NaOCl (1 mL per 100 seeds) for 11 minutes, followed by five 1-min rinses with sterile water. Seeds were then pre-germinated in 12×12 cm square plates (Greiner Bio-One, cat: 688161) containing Long-Ashton nutrient solution solidified with agar (Supplementary Table **14**). After two days of incubation in the dark, germinated seeds were transferred to fresh square plates. At the time the transfer, four seedlings per plate were inoculated by applying 0.5 mL of bacterial suspension or mock solution directly to the roots. Plates were allowed to dry for 30 min before sealing. Four replicate plates, each one containing four plants, were prepared for each condition.

For the growth-chamber pot assay, 0.3 L pots (7 cm x 7 cm x 7 cm) were fitted with a mesh at the bottom and filled with fine quartz sand sterilized by dry heat at 120°C for 3 hours. Three surface-sterilized seeds were placed equidistantly in each pot, and 1 mL of the bacterial suspension or mock solution was dispensed directly onto the seeds, which were then covered with approximately 1 cm of sand. Plants were sub-irrigated with Long-Ashton nutrient solution containing 32 µM phosphorus (Supplementary Table **14**). Four days after sowing, seedlings were thinned to one plant per pot. Ten pots were used per each treatment.

### 2.11. Statistical analysis

Each genotype’s response to the inoculum was calculated as follows (Equation 7):

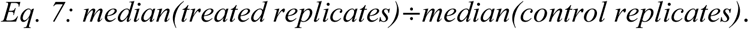

For Fig. **5**, the response of individual replicates to the inoculum was calculated as follows (Equation 8):

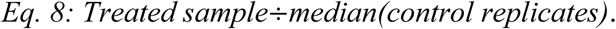

Differences between treated and control genotypes were assessed using the Kruskal-Wallis test, followed by Dunn’s *post hoc* test (fdr adjustment for multiple corrections). Plants were categorized as either ‘positive’ or ‘negative’ responders to the inoculum for each phenotypic trait analyzed. Specifically, if the ratio of the treated plants to the median of control plants for a given genotype exceeded 1.20, the genotype/plant was classified as a positive responder. Conversely, if the ratio fell below 0.80, it was categorized as a negative responder. Genotypes/plants with a ratio between 0.80 and 1.20 were considered ‘non-responding.’ This classification allowed us to identify three distinct conditions for phosphate accumulation and shoot biomass, which were used to account for differences in bacterial composition and diversity. The phyloseq (version 1.48.0), vegan, reshape2 (version 1.4.4), and ggplot2 (version 3.5.1) packages in R were utilized to generate stacked bar plots of bacterial communities and to perform both β-diversity analysis (using the Bray-Curtis distance matrix) and α-diversity analysis (Shannon and Observed indexes). Statistical differences in α-diversity between phenotypic response groups were assessed through the Kruskal-Wallis test, followed by Dunn’s *post hoc* test (fdr adjustment for multiple corrections). The Log2foldchange function from the DESeq2 package in R (version 1.44.0) with correction for multiple comparisons was used to examine differences in bacterial ASV presence between conditions of positive and negative phosphate and root biomass responses. Broad sense heritability for differentially abundant ASVs in leaf phosphate and root biomass variation was calculated in R with the Imer method from the Ime4 package (version 1.1.37). Pearson correlations from the *cor.test* function in the ggpubr package in R (version 0.6.0) were used to study correlations between the taxonomic ranks of fungal and bacterial communities.

## 3. Results

### 3.1 Physiological and nutritional variability across 128 *Lactuca sativa* accessions depend on genotypes

To broadly assess the impact of an inoculum containing arbuscular mycorrhizal fungi (AMF) and phosphate-solubilizing bacteria (PSB), 128 fully sequenced *Lactuca sativa* accessions were cultivated in pots under low-phosphate conditions, either with or without the inoculum treatment. Although all the plants were grown in a non-sterile soil-sand mixture, those inoculated showed a tendency toward higher mycorrhization rates compared with untreated plants (Fig. S**1**, Supplementary Table **15**). Plant responses were first evaluated across two key agricultural parameters: leaf-soluble phosphate concentration and shoot biomass variation. Results showed substantial variation in responses across genotypes, with many genotypes demonstrating positive responses - considered as 20% greater than non-responders-in both leaf soluble phosphate accumulation (25 genotypes, Fig. **1a**) and shoot biomass variation (28 genotypes, Fig. **1b**, Supplementary Tables **16,17**) consistently with growth curves obtained with phenospex data (Fig. S**2)**. Only 9 genotypes exhibited a decrease in leaf phosphate, while 30 genotypes showed a reduction in shoot biomass due to the treatment (Fig. **1a,b**). These findings highlight the strong intraspecific variation driving divergent responses among genotypes within the same plant species, an observation that is further reinforced by the high consistency in variation between treated versus mock replicates (Fig. **1a,b**, right side).

**Fig. 1.**
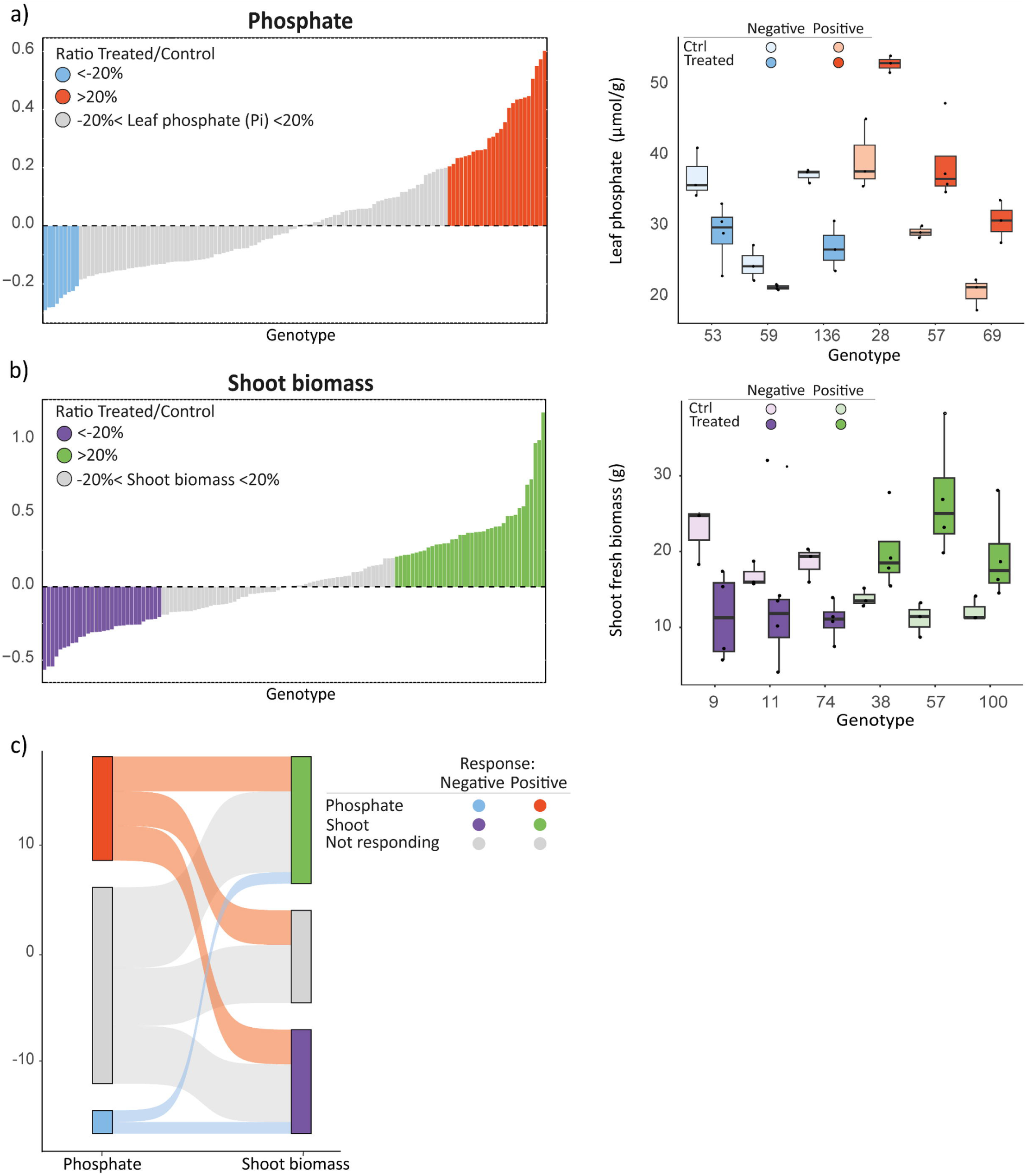
Leaf phosphate and shoot biomass variation in response to the microbial consortium. **a,b)** Bar plots depicting genotype responses in terms of leaf phosphate accumulation (**a**, left) and shoot biomass variation (**b**, left). In panel A, Y-axis values are expressed as median [Pi]treated / median [Pi]control, while in panel B, they represent median treated shoot biomass (g) / median control biomass (g). Genotypes with a 20% or greater increase in response are highlighted in red **(a)** and green **(b)**, whereas those with a 20% or greater decrease are shown in blue **(a)** and purple **(b)**, genotypes within the middle range are represented in gray. The boxplots on the right side of panels A and B display representative genotype responses for phosphate (**a**, right) and shoot biomass (**b**, right) using the same color scheme for positive (red/green) and negative (blue/purple) responses. **(c)** A Sankey diagram illustrates genotype response shifts due to the inoculum for phosphate accumulation (left) and shoot biomass variation (right), following the same color coding.

Interestingly, the fluctuations of these two parameters appeared independent of each other (Fig. **1c**). Specifically, among the 25 plants that accumulated more phosphate, 6 showed an increase in shoot biomass, 6 showed a decrease, and 13 exhibited no change. Similarly, among the 6 plants that accumulated less phosphate, 4 displayed an increase in biomass, while 2 showed a decrease (Fig. **2a**). By considering all measured morphological and physiological plant parameters in response to the treatment, we identified two main groups of response (Fig. **2a**). The analysis revealed a strong correlation between spectral parameters, such as hue and greenness, and various physiological indicators, including NCPI, PSRI, and NDVI. In contrast, morphology-related traits, such as digital biomass, shoot and root fresh biomass, and leaf area, formed a distinct cluster characterized by a more uniform response pattern. Interestingly, while some physiological traits (spectral parameters like NCPI, PSRI, and NDVI) were strongly correlated, the morphological response was highly variable and often uncoupled, as for the case of shoot biomass and phosphate accumulation mentioned above. Along the y-axis, three distinct response clusters emerged, independent of plant variety.

**Fig. 2.**
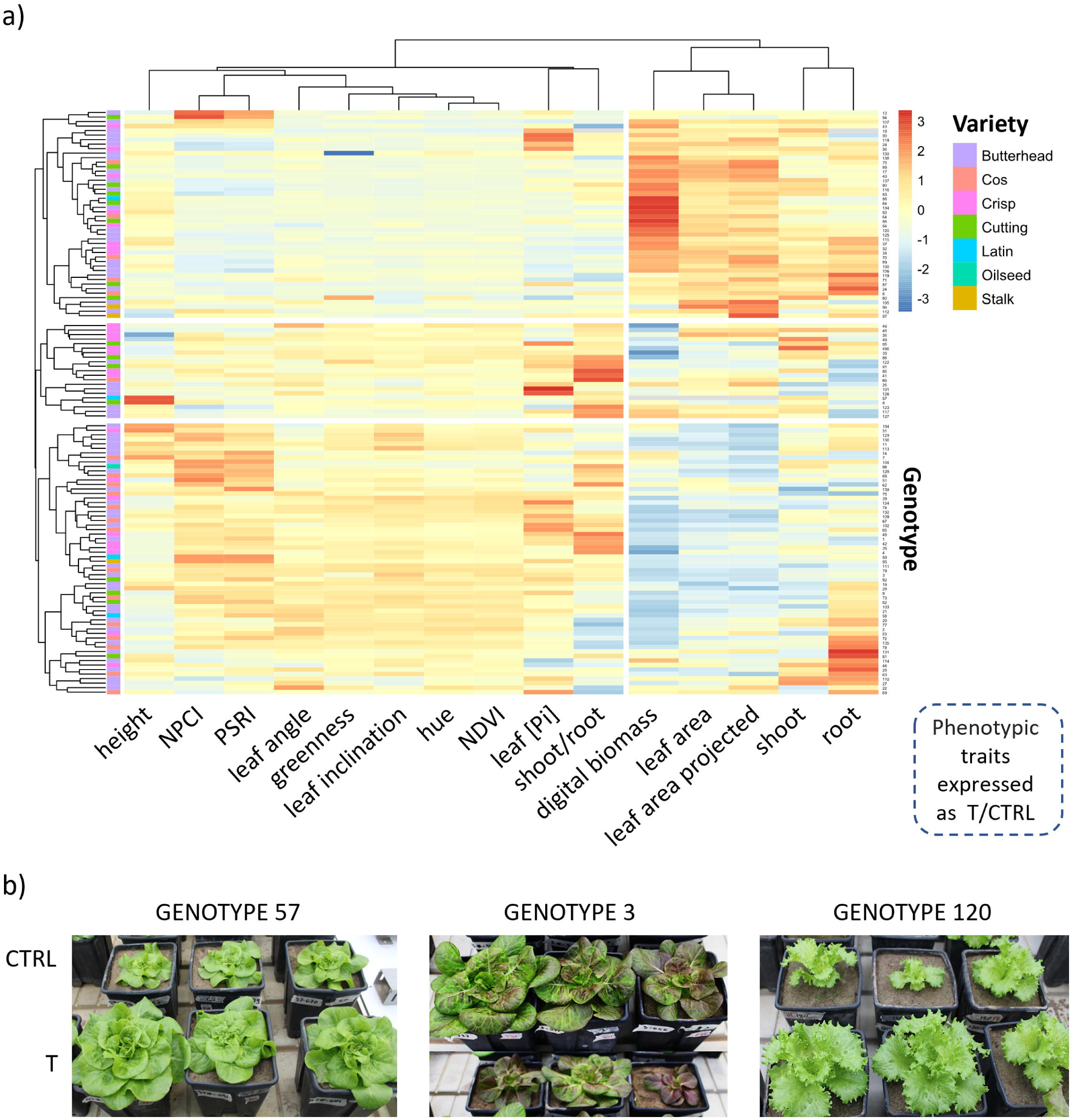
Overall phenotypic and physiological plant responses to the soil microbial inoculum. **a)** Hierarchical clustered heatmap of z-scaled genotype responses, expressed as median(treated)/median(control). High values are shown in red, low values in blue, and intermediate values in white. A filled rectangular shape on the left indicates the varieties to which the genotypes belong. **b)** Images of three representative genotypes, with control plants above and treated plants below.

For some genotypes, phenotypic variation resulting from inoculation was visibly apparent by the end of the experiment, as shown in Fig. **2b** and **S2**. Notably, genotype 57 stood out as a positive responder due to the pronounced phenotypic differences between mock and treated plants, coupled with a significant increase in leaf-soluble phosphate (Dunn *post-hoc*, p-value=0.034), as highlighted in Fig. **1a**.

Together, these results suggest that lettuce genetic variation plays a key role in shaping the outcomes of beneficial plant-microbe interactions at the soil-root interface.

### 3.2 Phosphate and shoot biomass responses to the microbial inoculum are linked to three SNPs in the lettuce genome

To better understand how genetic variation influences plant responses to inoculation, we conducted a GWAS to explore potential associations between lettuce SNPs and various phenotypic traits (Fig. **3**, S**3**, S**4**). Due to high correlations among some traits, four phenotypic parameters (Leaf area index, leaf area projected, light penetration depth, and maximum height) were excluded to streamline the dataset (Fig. S**3a,b)**.

**Fig. 3.**
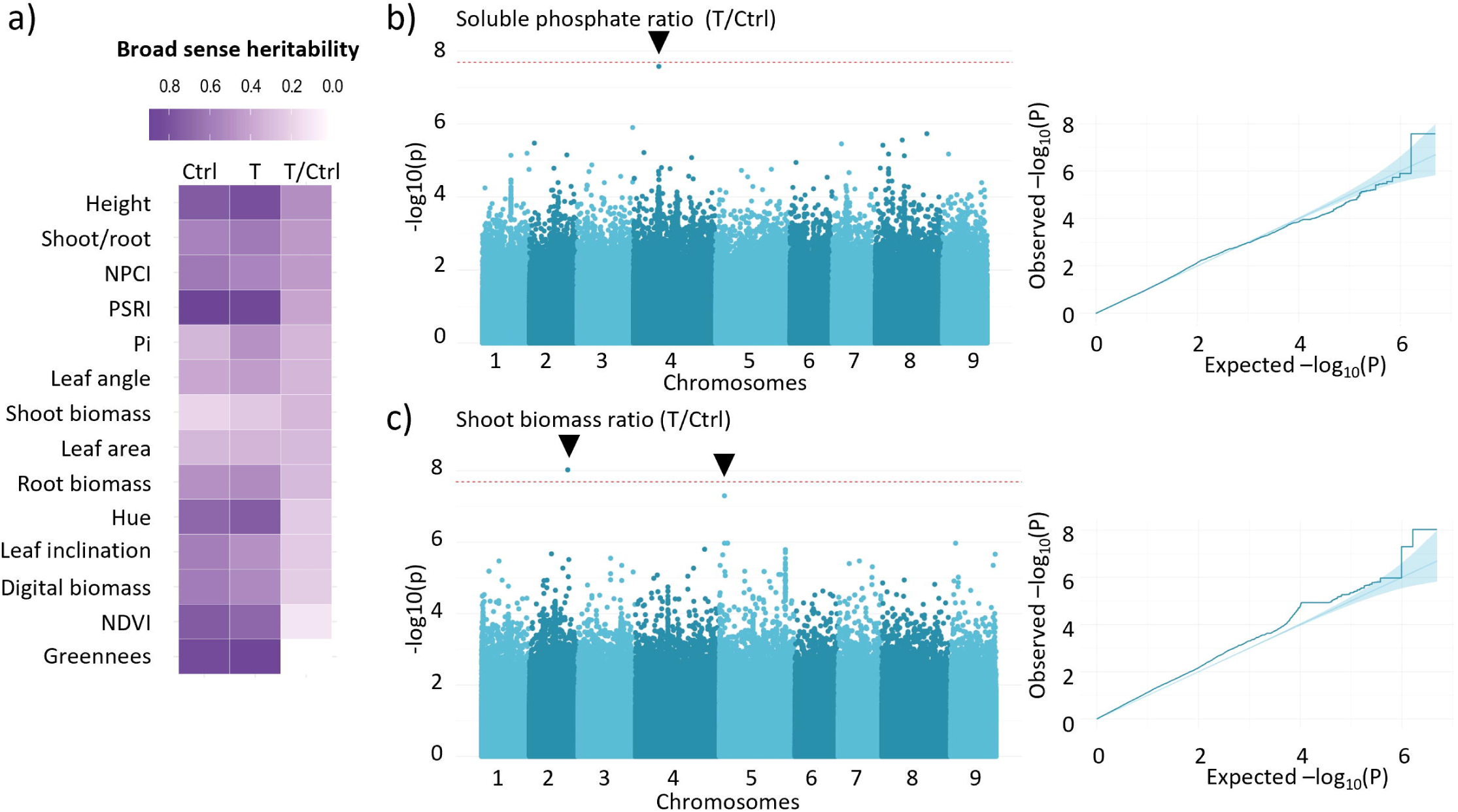
Genetic basis of the nutritional and physiological response of 131 *Lactuca sativa* genotypes to a microbial consortium. **a)** Heat Map displaying broad-sense heritability values calculated for control (Ctrl) and treated (T) plants, as well as their ratio (T/Ctrl). **b,c)** On the left hand, Manhattan plots show the correlation between lettuce SNPs and variations in **(b)** leaf-soluble phosphate ratio and **(c)** shoot biomass ratio. On the right, QQ plots illustrate expected false positive p-values versus observed values.

**Fig. 4.**
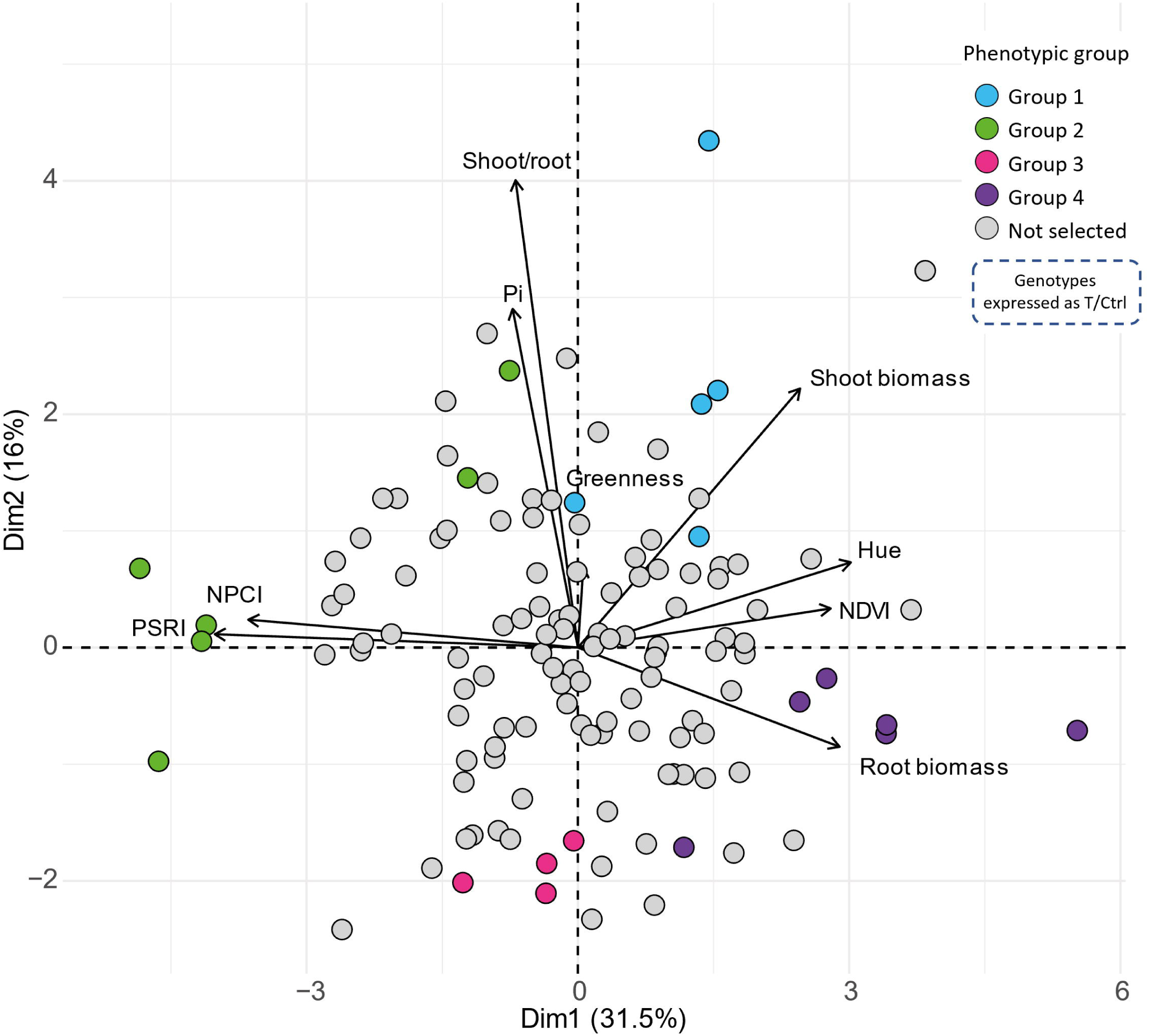
Multiple phenotypes integration disentangles plant response to the microbial inoculum. Principal component analysis of genotype responses to inoculum treatment, expressed as median (treated values)/median (control values). Genotypes chosen for metabolite quantification, representing the four distinct phenotypic groups, are color-coded as blue, green, pink, and purple. Non-selected genotypes are shown in gray.

We calculated broad-sense heritability (H^2^) for traits of treated plants, control plants, and their ratio (reflecting the response to inoculation, Fig. **3a**, Supplementary Table **18**). PSRI had the highest heritability for both control (H^2^ = 89.62%) and treated (H^2^ = 87.18%) plants, while shoot biomass had the lowest heritability for control (H^2^ = 19.05%) and treated (H^2^ = 24.67%) plants. Overall, heritability decreased for plant response (ratio) compared to the control and treated conditions. The highest H^2^ for plant response was observed for head height (H^2^ = 50.97%), while the lowest was for greenness (H^2^ = ∼0). Notably, shoot biomass variation (T/Ctrl, H^2^ = 32.56%) and soluble leaf phosphate variation (T/Ctrl, H^2^ = 33%) were among the few traits where H^2^ increased compared to both control and treated plants (Fig. **3a**).

For soluble leaf phosphate, we identified a highly significant SNP on chromosome 4 (position: 117030769, allele frequency = 32.3%, p.wald = 6.79e-08, Fig. **3b**). Additionally, two SNPs, one on chromosome 2 (position: 22753367, allele frequency = 8.3%) and another on chromosome 5 (position:172536404, allele frequency = 49.2%), were significantly associated with shoot biomass variation (p.wald = 2.56e-08 and 1.43e-08, respectively, Fig. **3c**, Table **1**). For other traits, GWAS was performed on the response (T/Ctrl) parameters with H^2^ greater than 20% (Fig. S**4**). This analysis showed no significant SNP associations for the shoot/root ratio (Fig. S**4a)**, leaf angle ratio (Fig. S**4e)**, and root ratio (Fig. S**4f)**. However, several significant SNPs associated with NPCI (Fig. S**4c)** and PSRI (Fig. S**4d)** were identified. On the other hand, head height (Fig. S**4b)** displayed highly polygenic patterns. Table **1** provides a comprehensive list of all significant SNPs identified in this analysis, with further details and raw data reported in Supplementary Table **19** available at Zenodo (DOI 10.5281/zenodo.14224764).

**Table 1.** List of SNPs significantly associated with phenotypic and physiological traits.

These findings highlight the complex genetic architecture underlying lettuce’s response to inoculation.

### 3.3 Integrating plant phenotypic and metabolic traits allows a detailed clustering of inoculum effects on plants

To identify clusters of genotypes that showed an overall similar plant phenotypic response to the inoculum, we conducted principal component analysis (PCA) on the trait ratios for each genotype (T/Ctrl, Fig. **4**, Fig. S**5**). Given the strong correlations among certain phenotypic traits (Fig. S**5a**), we used only a few independent and highly descriptive ones to define the phenotypic space of the genotypes (Fig. **4**). This approach effectively separated physiological parameters (NPCI, PSRI, Hue, NDVI) from morphological ones (shoot and root biomass and their ratio). Notably, the leaf phosphate vector is orthogonal to shoot biomass, nearly overlapping with the shoot/root ratio, while the phosphate and root vectors point in opposite directions, reflecting plant physiological responses to phosphorus availability or deficiency. In summary, this method identified four distinct response groups across the lettuce population capturing the plant phenotypic variation in response to microbial inoculation. Among these clusters, we selected 28 genotypes to further analyze their leaf metabolites (Fig. S**5b**), which confirmed the initial classification and led to the exclusion of six genotypes that did not fit into any group. The remaining 22 genotypes, represented by 91 plants occupying diverse plant phenotypic space, were used for downstream analysis, with the hypothesis that plants with similar responses to the inoculum exhibit analogous microbial interactions.

### 3.4 The four phenotypic response groups are associated with diverse root-associated microbial communities

To explore the relationship between plant phenotypic responses and root microbial communities, we analyzed the fungal and bacterial community composition across all substrates that were used in our experimental setup. The growing medium (sand and water) exhibited the highest bacterial Shannon and Observed diversity, while roots displayed the lowest diversity, followed by germination soil and microbial inoculum (Fig. S**6a,b,c,** Fig. S**7a,b,c)**. Although the inoculum’s richness was similar to or lower than that of the soils, it enhanced the Shannon diversity index in treated plants compared to controls (Fig. S**6a,b,** Fig. S**7a,b,c**). Additionally, each compartment had a distinct bacterial composition (Fig. S**6d,** Fig. S**7d**). The dominant bacterial phyla across all compartments were Proteobacteria and Actinobacteriota, while for fungi Ascomycota predominated (Supplementary Table **20a,b**). The inoculum showed enrichment in Firmicutes (12% in inoculum compared to 5% in roots, 2% in growth soil, and 0% in germination soil) and Bacteroidota (6% in inoculum, 5% in roots, 4% in growth soil, and 3% in germination soil) (Supplementary Table **20a**). Notably, an amplicon sequence variant corresponding to *Chryseobacterium idoltheticum* was detected in plant roots and assigned to ASV1713 (Supplementary Table **21**). The Glomeromycota phylum, to which *Rhizophagus irregularis* and *Funneliformis mosseae* belong, was present in the roots and in the inoculum but not in the other compartments (Supplementary Table **20b**).

To achieve greater resolution in individual plant responses, we examined root-associated bacterial diversity at a single-replicate level, as described in the Materials and Methods (Eq. **8**) and Supplementary Table **10**. This analysis showed statistically significant differences in fungal β-diversity (PERMANOVA, p-value = 0.018, Fig. **5a**) among the four previously identified phenotypic groups, but not in their α-diversity (Fig. S**8a**). For bacteria, instead, no significant differences were observed in the bacterial Shannon and Observed α-diversity indices among the four previously identified phenotypic groups (Fig. S**8b**).

**Fig. 5.**
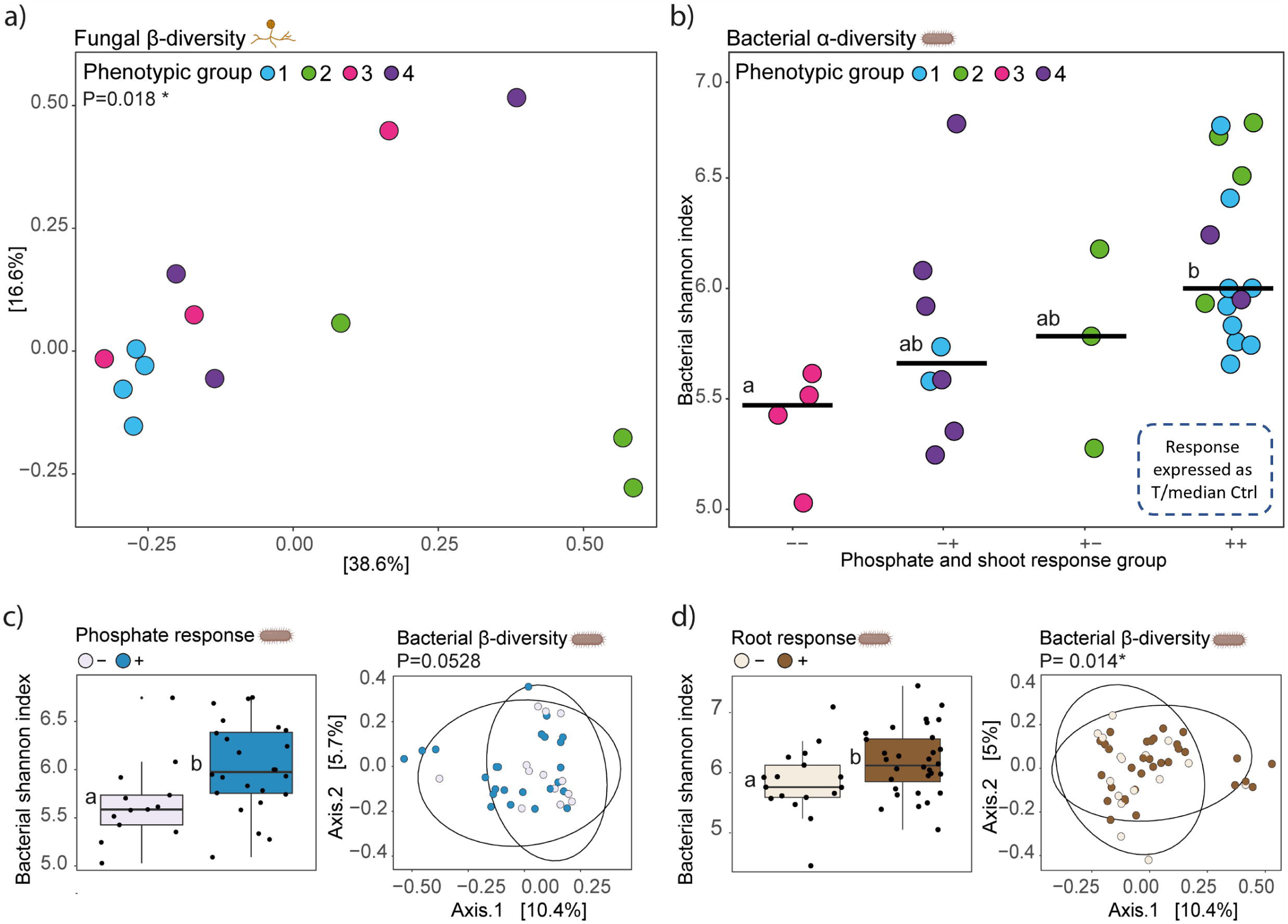
Root-associated microbial variation at the base of divergent plant phosphate and shoot responses. **a)** Fungal Bray-Curtis dissimilarity distance matrix between phenotypic groups (highlighted by colors). **b)** Bacterial Shannon index variation across the four response intervals: “--” denotes negative responders, “++” positive responders, with “-+” and “+-” representing intermediate responses for phosphate and shoot biomass variation, respectively.. Colors indicate the phenotypic group for each replicate. **c)** Bacterial Shannon index and Bray-Curtis distance matrix variation comparing plants with low (–, in gray) and high (+, in blue) leaf phosphate accumulation. **d)** Shannon index and Bray-Curtis distance matrix variation comparing plants with negative (–, in light brown) and positive (+, in brown) leaf phosphate accumulation.

Building on this result, we narrowed the analysis to phosphate and shoot response and observed that bacterial α- diversity indices increased with progressively higher leaf phosphate and shoot biomass response (Fig. **5b**). This trend was most pronounced between plants exhibiting completely negative (−−) versus completely positive (++) responses (Dunn post hoc, adjusted p-value = 0.014). In contrast, no comparable differences were observed for fungi (Fig. S**9a,b**). To determine the main driver of this correlation, we further separated the response into phosphate and shoot components. Fungal diversity did not consistently vary with leaf phosphate (Fig S**9c**), shoot biomass (Fig. S**9d**), root biomass (Fig. S**9e**, right part) and shoot/root biomass responses (Fig. S**9f**). The only notable exception was a larger variation in root biomass, which correlated with a significantly higher fungal Shannon index (Dunn post hoc, adjusted p-value=0.0098, Fig. S**9e**).

In bacteria, leaf phosphate variation was significantly associated with a distinctively richer bacterial community (Dunn post hoc, adjusted p-value = 0.012; Fig. **5c**, Fig. S**10a,b**) and showed a near-significant effect on bacterial β- diversity (PERMANOVA, p-value = 0.0528; Fig. **5c**). In contrast, shoot biomass variation was not associated with changes in α- or β-diversity (Fig. S**10c**). Among the other morphological parameters, only root variation was linked to changes in both bacterial α- and β- diversity (Fig. **5d**). Specifically, plants with larger root systems hosted significantly richer bacterial communities for both Shannon (Dunn post hoc, adjusted p-value = 0.036, Fig. **5d**) and Observed indices (Dunn post hoc, adjusted p-value = 0.025, Fig. S**10d**) compared to those with smaller roots. The bacterial diversity between the two conditions was also validated by β-diversity, (PERMANOVA, p-value = 0.014, Fig. **5d**). Conversely, the shoot/root biomass ratio was not linked to shifts in bacterial β- and α- diversity (Fig. S**10e**). The positive association between bacterial α- diversity and leaf phosphate variation was further confirmed in a subsequent experiment, in which plants were treated with the same synthetic community, in controlled growth chamber condition, and with the AM fungus alone (Fig. S**11**, Supplementary Table **22,** Supplementary Material and Methods).

Together, these findings underscore the tight regulation of root-associated fungal and bacterial diversity in relation to key plant nutritional parameters, such as leaf phosphate accumulation, while also highlighting its associations with morphological changes, particularly in root system development.

### 3.5 Positively responding plants show enrichment of differentially abundant ASVs from Burkholderiales and Flavobacteriales

To identify the ASVs that are differentially abundant between plants with positive and negative responses in terms of phosphate accumulation and root biomass, at the base of the observed differences in diversity, we performed a Log2 fold-change analysis. This approach revealed 22 bacterial ASVs that were significantly more abundant (adjusted p-value < 0.05) in plants accumulating higher phosphate levels in response to the inoculum, with no ASVs significantly enriched in the opposite condition (Fig. **6a**). In addition, 6 bacterial ASVs showing a base mean above 10, and 16 ASV with a base mean below 10, were enriched in plants exhibiting a positive root biomass response to the treatment (FIG. **6b**, Fig. S**12a**).

**Fig. 6.**
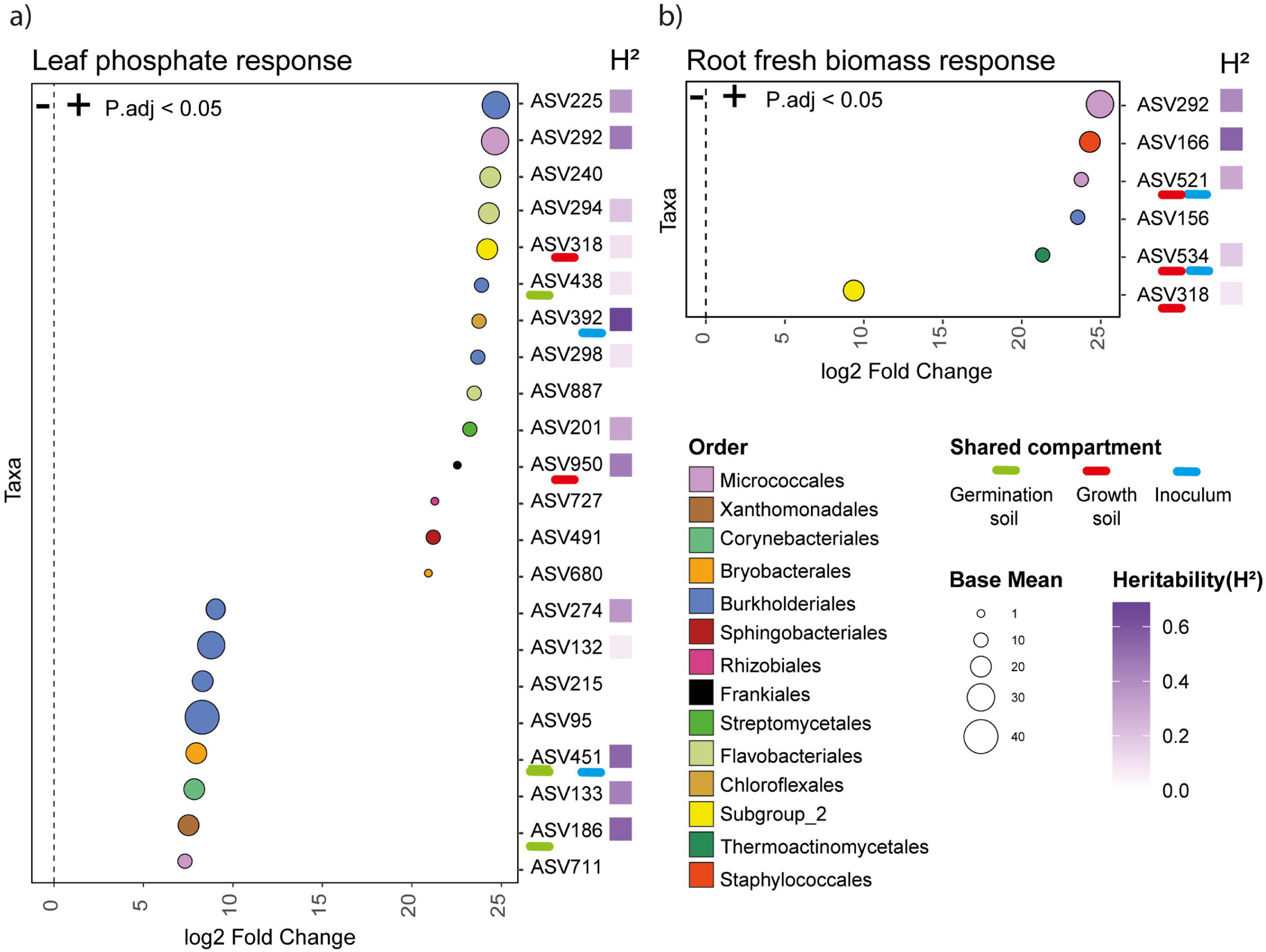
Differential abundance of ASVs in plants with positive and negative leaf phosphate accumulation and shoot fresh biomass response. **a)** Log2 fold-change values representing ASVs that are more abundant in plants with high (+) and low (−) phosphate accumulation in response to the inoculum. **b)** Log2 fold-change values representing ASVs with a base mean above 10 that are more abundant in plants with a positive (+) and negative (−) root fresh biomass variation in response to the inoculum. Each dot represents an ASV, with colors indicating its taxonomic order and size representing the base mean effect. ASVs shared with the germination soil are underlined in green, in the growth soil in red and in the inoculum in blue. Broad-sense heritability (H²) is shown using a purple color scale, with higher H² values in dark purple and lower values in lighter shades.

To determine if these ASVs were shared with other compartments and speculate a possible origin, we found that, for phosphate response, ASVs 438 and 186 were uniquely associated with the germination soil, ASVs 950 and 318 were specific to the growth soil, and ASV392 were uniquely shared with the inoculum. In the roots, we identified two ASVs (2775 and 1037) that were uniquely shared with the germination soil and the inoculum, respectively. Additionally, two other ASVs were shared between both the growth soil and the inoculum.

Almost half of the ASVs enriched in the positively responding plants belong to Burkholderiales (7 out of 22) and Flavobacteriales order (3 out of 22, Fig. **6a**). In the root biomass response, the Burkholderiales represents 3 out of the 22 total ASVs, while only one Flavobacteriales ASVs is present (Fig. **6b**, Fig. S**12a**).

To evaluate how plant genotype contribute to variation in the presence of differentially abundant ASVs under contrasting conditions, we calculated broad-sense heritability (Supplementary Table **23**). Eight ASVs, including two from the order Burkholderiales (ASVs 225 and 274), exhibited H² values greater than 0.30 in response to phosphate variation. Among these, ASVs 392 and 451 showed the highest heritability (H²= 0.69 and 0.54, respectively), and were also detected in the microbial inoculum (Fig. **6a**). For root biomass variation, five ASVs exhibited a H² values above 0.30, with ASV 521 notably shared with the microbial inoculum (Fig. **6b**, Fig. **S12a**).

To explore any possible link between the fungal community (with a particular focus on the AMF used in this experiment) and the bacterial ASVs enriched under conditions of increased leaf phosphate and root biomass, we performed correlation analyses between the two communities with a particular focus on Burkholderiales that represented the most enriched order. We found a positive correlation (Pearson’s correlation, R=0.39, p-value=0.0051) between the root-associated Burkholderiales and Glomeromycota relative abundances (Fig. S**12b**). After analyzing each differentially enriched ASV in the two conditions described above, ASV292 (Micrococcales order) showed a positive correlation with the relative abundance of the Glomeromycota phylum in both conditions of positive phosphate and root biomass response (Fig. S**12c,d**). Interestingly, ASV292 had among the highest heritability values in both the two conditions (H²=0.46). Other significant correlations were detected between ASV294 (Flavobacteriales order), ASV186 (Xanthomonadales order) and ASV438 (Burkholderiales order) and the Glomeromycota relative abundance in positive responding plants for leaf phosphate (Fig. S**12c**). Additionally, another member of Micrococcales order, ASV521, was correlated with the fungal community in plants with a positive root biomass response (Fig. S**12d**).

In summary, we identified 44 bacterial ASVs associated with increased phosphate accumulation and improved shoot biomass response. Among these, five ASVs were positively correlated with the Glomeromycota fungal phylum, suggesting a possible effect of fungal presence on the bacterial community and, consequently, on plant nutrient acquisition.

### 3.6 Effects of six selected ASV-corresponding isolates on leaf phosphate content and shoot biomass under plate and pot conditions

To test causal relationships between the identified ASVs, plant biomass, and leaf phosphate concentration, we selected six isolates from the lettuce-root culture collection whose sequences were highly similar (>97% identity) to ASVs that were significantly enriched in our dataset. These isolates were used in both plate assays and pot experiments, as described in the Materials and Methods section (Section 2.10), to assess phosphate-related phenotypes with and without bacterial inoculation.

In the plate experiment, the isolate corresponding to ASV438 (Comamonadaceae_1), significantly increased leaf phosphate content in genotype Butterhead 38. Similarly, the isolate corresponding to ASV225 (Comamonadaceae_2), increased leaf phosphate content in genotype Cos 100 compared with both the mock treatment and the synthetic community composed of all six isolates (Fig. **7A**). None of the tested isolates, either individually or in combination, significantly affected shoot biomass, measured as fresh weight, under plate conditions, except for the Xanthomonadaceae isolate, corresponding to ASV186, which reduced shoot biomass in genotype 100 compared with all other treatments (Fig. **7A**). Under pot conditions, the synthetic community composed of all six isolates significantly increased fresh plant biomass in two of the three genotypes tested, namely genotypes Butterhead 24 and 38 (Fig. **7B**).

**Fig. 7.**
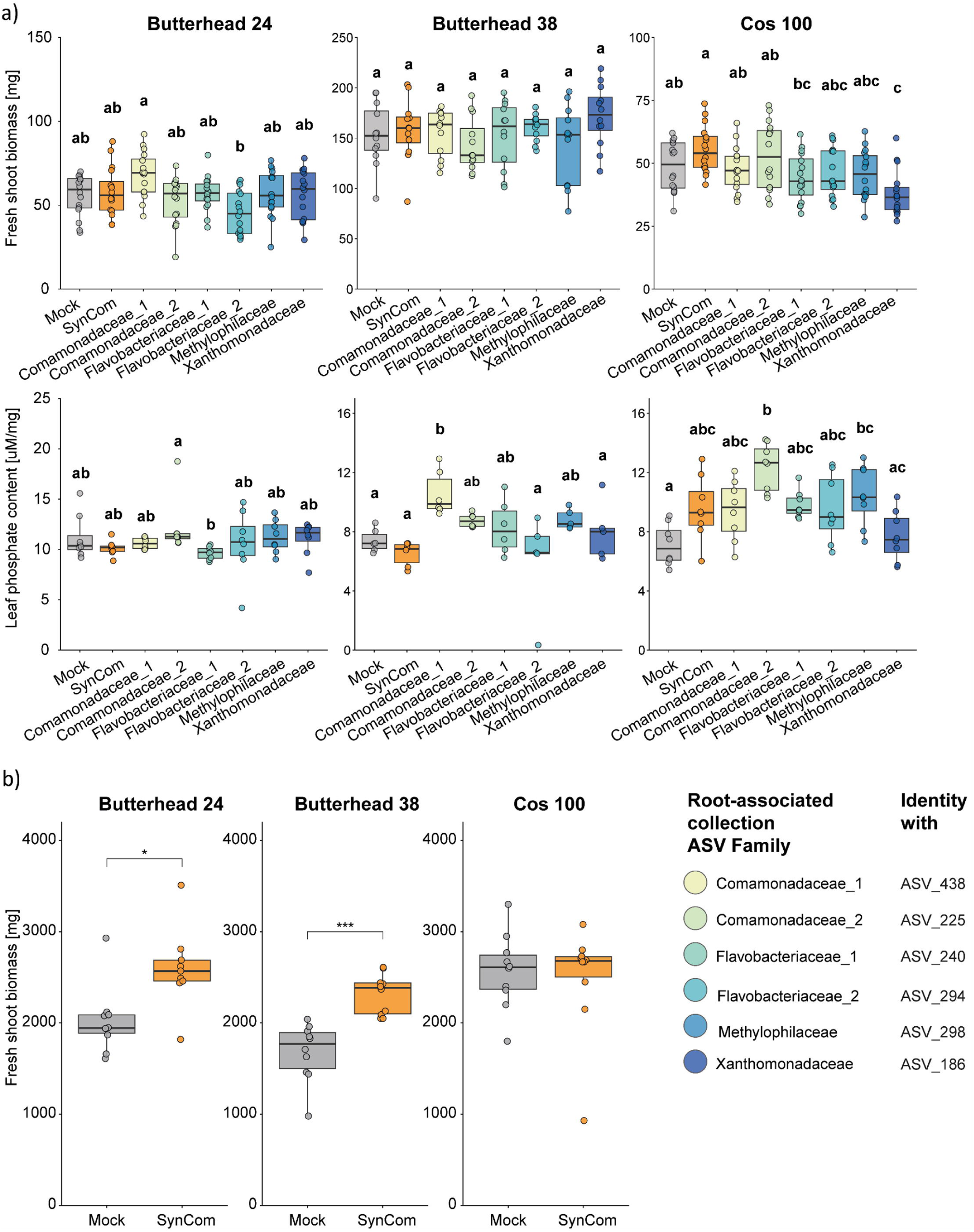
Effects of beneficial ASVs and their combination on plant shoot biomass and leaf phosphate concentration under plate and pot conditions. **(a)** Effects of individual bacterial ASVs (blue shaded boxplots, grouped at the family level) and their combined synthetic community (SynCom, orange) compared with mock-treated plants (gray) on shoot fresh biomass (upper panels) and leaf phosphate content (lower panels) in genotypes 24, 38, and 100 under plate conditions. **(b)** Effect of the SynCom on shoot fresh biomass in genotypes 24, 38, and 100 under pot conditions.

Together, these experiments provide evidence for a causal relationship between ASVs enriched in plants showing positive responses to the treatment and plant phenotypes following inoculation with corresponding bacterial isolates. Individual isolates increased leaf phosphate content in a genotype-dependent manner, although they did not produce a consistent effect on shoot biomass *in vitro*. However, their combined application as a synthetic community under pot conditions significantly increased plant biomass in two of the three tested genotypes.

## 4. Discussion

It is increasingly evident that plant intraspecific variation is crucial in shaping the plant microbiota and plant responses to it. However, the specific genetic regions responsible remain largely unknown and it is not clear if the same genomic variants only govern interactions with specific microorganisms or a broad spectrum of them (Peiffer *et al*., 2013; Bulgarelli *et al*., 2013b; Hassani *et al*., 2018; Deng *et al*., 2021).

In this study, we explored the connection between genetic diversity and plant responses to a soil microbial inoculum by conducting a large-scale greenhouse experiment on a population of 128 *Lactuca sativa* genotypes. Our findings revealed a strong influence of genetic diversity on both the phenotypic and physiological responses of plants. Specifically, we observed that different genotypes exhibit varied responses to the treatment both in morphological traits, such as root and shoot biomass, leaf angle, and in physiological traits, such as phosphate accumulation.

Supporting these findings, a GWAS analysis identified specific genetic regions in the lettuce genome associated with responses to microbial inoculation. For key agronomic traits such as leaf phosphate concentration and shoot biomass, we identified one SNP on chromosome 4 and two SNPs on chromosomes 2 and 5, respectively, that are linked to variations resulting from the treatment. Overall, heritability values decreased when evaluating the plant’s response to microbial inoculation compared to the median values observed in treated and untreated plants. However, for the two traits mentioned above, leaf phosphate concentration and shoot biomass, heritability increased in response to the inoculation. This underscores the genetic control exerted by plants on key parameters that are influenced by the interactions with beneficial microorganisms. Identifying the putative genes involved in establishing interactions that determine these beneficial fitness phenotypes in the host will be crucial and will require further investigation using transcriptomic and metabolomics approaches, taking into account the large Linkage Disequilibrium in *Lactuca sativa*, where pairwise correlation coefficient dropped to half of its maximum value at 177.7 kb (Wei *et al*., 2021).

Beyond investigating the plant genetic base of differential plant responses to soil microorganisms, our results revealed a strong connection between the four identified plant phenotypic response groups and the root-associated fungal β- diversity. Conversely, root-associated bacterial communities were associated with the plant response through their α-diversity, showing increased Shannon and Observed indexes with progressively more positive shoot biomass and leaf phosphate accumulation. These findings suggest a potential link between plant phenotypic responses to microbial inoculum, phosphate accumulation and, the richness and diversity of root-associated microbial communities.

Furthermore, we identified 22 bacterial ASVs differentially enriched in plants with a positive leaf phosphate response to the inoculum and 22 ASVs enriched in plants with increased root biomass as a result of the treatment. Two of them, ASV392 (belonging to the Chloroflexales order) and ASV1037 (belonging to the Corynebacteriales order) for leaf phosphate and root biomass, respectively-were uniquely shared with the inoculum. Given that 7 out of 22 ASVs associated with improved leaf phosphate assimilation belong to the Burkholderiales order, known to be obligate endosymbionts of the cytoplasm or intracellular structures of AMF (Basiru *et al*., 2023; Zhang *et al*., 2024), we investigated their potential association with the AMF present in our inoculum, which belongs to the phylum Glomeromycota. Our findings revealed a positive correlation between root-associated Burkholderiales order and Glomeomycota phylum. Among Burkholderiales, ASV438, was positively correlated with Glomeromycota relative abundance in positive responding plants for phosphate accumulation. To establish causality between differentially enriched ASVs and their effects on plant phenotypes, as hypothesized on the basis of the bioinformatic analyses, we performed experimental validation using six bacterial strains isolated from lettuce roots. These strains were tested individually and in combination (SynCom) under both *in vitro* plate conditions and in sterile sand in growth chamber experiments, across three lettuce genotypes. Overall, these experiments demonstrated that the selected bacterial strains can significantly influence plant growth and leaf phosphate content; however, their effects are strongly genotype-dependent.

Overall, these findings highlight three key considerations for improving the efficacy of microbial inoculants when applied to soil. First, plant genotype is crucial and should be considered in the breeding process to select plants capable of establishing beneficial interactions with soil microorganisms, which can enhance agronomic parameters such as leaf phosphate content and shoot biomass (Escudero-Martinez & Bulgarelli, 2023). Second, selecting plant morphological traits, such as root biomass and leaf phosphate concentration, that promote the association with richer soil microbial communities can further enhance the functional benefits these microorganisms provide to plants. Thirdly, integrating genetic and phenotypic variation can facilitate the identification of bacterial strains linked to variation in agriculturally important phenotypic traits under different conditions.

Further research should target fungal communities using tailored experimental designs and increased sequencing depth to better understand their interactions with bacterial assemblages. In addition, given the challenges in tracking inoculated bacteria throughout the experiment (only *Chryseobacterium* was recovered) future efforts should prioritize methods to improve the detection and persistence assessment of introduced strains.

The results of this research not only clarify some of the mechanisms shaping interactions between soil microorganisms and *Lactuca sativa* roots but also provide a foundation for future breeding efforts. These should focus on selecting optimal combinations of genotypes, morphological traits, and microbial communities to develop resilient plants that sustainably meet the needs of a growing global population without compromising environmental health.

## ACKNOWLEDGMENTS

We gratefully acknowledge the reviewers’ comments, which have substantially improved the manuscript. We would like to express our gratitude to Sofìa Cristina Somoza, Niccolò Forin, Nadine De Biasio, Enrico Cortese, Gregory Saccozza, Filippo Bomitali, Solomon Simon, and Michele Cappellari for their assistance with plant design preparation and sample collection. We also thank Andrea Mattarei for the ultra-analytical balance and Matteo Chialva for tips on data analysis. Finally, we are grateful to Lorella Navazio and Barbara Baldan for sharing their workspace. This work was supported by grants from the European Union (NextGenerationEU, Mission 4, Component 1, CUPD53D23022080001 PRIN-PNRR prot. P2022WL8TS to MG), the Ministry of the University and Research (PON Ricerca e Innovazione 2014-2020 PhD fellowship DOT1471523 to Ac) and the University of Torino (GTI grant to MG).

## AUTHOR CONTRIBUTION

ACa, MG and FV planned and designed the research. ACa, MG, CS, SS, FV, ACi and AA performed experiments and conducted fieldwork. ACa and PS conducted the phosphate and metabolite quantification. ACa performed DNA extraction. ACa, MG and PC analyzed data. CS, SS, AA, FV and MG supervised the work. AC, MG wrote the manuscript.

## DATA AVAILABILITY STATEMENTS

The datasets generated during the current study are available in the NCBI SRA repository as Bioproject PRJNA1215347. The R script used to obtain the results and the figures presented in this paper is reported as Supplementary script.

## The following Supporting Information is available for this article

**Fig. S1. Frequency of mycorrhization (F%) in control and treated samples.**

**Fig. S2. Digital biomass trends of genotypes with differential responses (9-11-74-38-57-100) over the course of the experiment.** Genotypes that show a decrease in biomass in response to the treatment are highlighted in purple, while those that experience an increase in biomass due to inoculation are shown in green.

**Fig. S3. Correlation matrix among all parameters measured by the Phenospex system. a)** Correlation matrix including all measured values, and **b)** correlation matrix showing differences between treated and control samples. Positive correlations are colored in red, negative correlations are in blue, and no interactions are in white. Dot size indicates the strength of the correlation, with larger dots representing stronger interactions.

**Fig. S4. GWAS of phenospex parameters**. **A-J**) Manhattan plots and QQ plots showing the correlation between SNPs in the *Lactuca sativa* genome and the following traits: **a)** shoot/root ratio, **b)** head height ratio, **c)** NPCI ratio, **d)** PSRI ratio, **e)** leaf angle ratio, **f)** root ratio, **g)** PSRI treated, **h)** height control, **i)** height treated, **j)** PSRI control.

**Fig. S5. Metabolite quantification to validate genotype phenotypic categorization**. **a)** Correlation Matrix among all parameters measured by the Phenospex system and the quantified metabolites. Positive correlations are colored in red, negative correlations are in blue, and no interactions are in white. Dot size indicates the strength of the correlation, with larger dots representing stronger interactions. **b)** Principal component analysis of responses to the inoculum in 28 selected genotypes, based on plant phenotypes and leaf metabolites. Genotypes chosen for root-DNA extraction, representing four distinct phenotypic groups, are color-coded in blue, green, pink, and purple. Non-selected genotypes are displayed in gray.

**Fig. S6. Fungal community richness and composition across different compartments.** Panels a,b display Shannon **(a)** and Observed **(b)** α-diversity indices across germination soil, growth soil, inoculum, and root samples. The lower section of each panel compares Shannon **(a)** and Observed **(b)** α-diversity between control (gray) and treated (brown) root samples. **c)** Rarefaction curves following size-based rarefaction across germination soil, growth soil, inoculum, and root samples. **d)** Bacterial community composition at the phylum level across compartments.

**Fig. S7. Bacterial community richness and composition across different compartments.** Panels a,b display Shannon **(a)** and Observed **(b)** α-diversity indices across germination soil, growth soil, inoculum, and root samples. The lower section of each panel compares Shannon **(a)** and Observed **(b)** α-diversity between control (gray) and treated (brown) root samples. **c)** Rarefaction curves following size-based rarefaction across germination soil, growth soil, inoculum, and root samples. **d)** Bacterial community composition at the phylum level across compartments.

**Fig. S8. Root-associated microbial variation at the base of phenotypic plant response**. **a)** Fungal and **b)** Bacterial Shannon (above) and Observed (below) α- diversity indices across the four phenotypic groups, indicated by different colors.

**Fig. S9.** Root-associated Fungal community variation at the base of divergent responses**. a)** Fungal Shannon and **b)** Observed α- diversity indices across the three response groups (−−,− +,++) for leaf phosphate and shoot biomass variation, with each replicate color-coded by the phenotypic group. c) Fungal Observed α-diversity diversity index comparing plants with positive (blue) and negative (gray) responses to leaf phosphate accumulation. **c-f)** Fungal α-diversity (Shannon, and Observed indexes) and β- diversity indices comparing positive and negative responses in leaf phosphate accumulation **(c)**, shoot biomass **(d)**, root biomass **(e)**, and shoot/root biomass variation **(f)**.

**Fig. S10. Root-associated bacterial community variation at the base of divergent responses**. **a)** Bacterial Observed α-diversity indices across the four response groups (−−,−+,+-,++) for leaf phosphate and shoot biomass variation, with each replicate color-coded by the phenotypic group. **b)** Shannon diversity index comparing plants with positive (blue) and negative (gray) responses to leaf phosphate accumulation. **c-f)** Bacterial β-diversity, Shannon, and Observed α-diversity indices comparing positive and negative responses in shoot biomass **(c)**, root biomass **(d)**, and shoot/root biomass variation **(e)**.

**Fig. S11. Leaf Pi response to fungal (Myc) and synthetic microbial community (SynCom) treatments under Pi deficiency. a)** Difference in soluble phosphate levels between mock-treated plants (gray), plants treated with AM fungus alone (Myc, light brown), and plants treated with the microbial inoculum (SynCom, dark brown). **b)** Shannon (top) and Observed (bottom) bacterial α- diversity indices comparing plants with strong (blue) or low (gray) leaf phosphate accumulation compared to mock-treated plants.

**Fig. S12. Correlation analysis between the root-associated fungal community and the bacterial ASVs enriched in the plants with a positive leaf phosphate and root biomass response. a)** Log2 fold-change values for ASVs, including those with a base mean below 10, showing their abundance in plants with positive (+) and negative (−) variations in root fresh biomass in response to the inoculum. Each dot represents an ASV, with colors indicating its taxonomic order and dot size representing the base mean effect. ASVs shared with germination soil are highlighted in green, those shared with growth soil in red, and those shared with the inoculum in blue. Broad-sense heritability (H²) is shown using a purple color scale, with higher H² values in dark purple and lower values in lighter shades. **b)** Pearson’s correlation between the relative abundance of Burkholderiales (on the x-axis) and the relative abundance of Glomeromycota (on the y-axis). **c-d)** Pearson’s correlation between the relative abundance of ASVs 292, 294, 186, 438, and 521 (x-axis) and the relative abundance of Glomeromycota (y-axis). The color of the confidence interval indicates the taxonomic order of each ASV. Panel **c** shows ASVs enriched in plants with a positive phosphate uptake response, while panel **d** shows those enriched in plants with increased root biomass.. Only statistically significant correlations are shown.

## Supplementary Tables

**Supplementary Table 1:** Preliminary lettuce phosphate starvation trial conducted prior to the large-scale experiment.

**Supplementary Table 2:** Greenhouse experimental design

**Supplementary Table 3:** Fertirrigation and inoculum administration plan

**Supplementary Table 4:** PlantEye scanning order

**Supplementary Table 5:** PlantEye measure per genotype

**Supplementary Table 6:** Fresh shoot and root biomass per genotype

**Supplementary Table 7:** Sampling design

**Supplementary Table 8:** Soluble leaf phosphate (Pi) quantification

**Supplementary Table 9:** Leaf metabolite quantification

**Supplementary Table 10:** Genotypes selected for DNA extraction

**Supplementary Table 11:** Sample information and quantification values for the genotypes selected for 16S metagenomic analysis

**Supplementary Table 12:** Reads remaining at each step of sample contamination removal

**Supplementary Table 13:** Percentage identity between the root-associated collection and the ASVs identified in this study.

**Supplementary Table 14:** Long Ashton recipe

**Supplementary Table 15:** Trouvelot assay data

**Supplementary Table 16:** Dunn post hoc test for leaf phosphate concentration between control and treated plants

**Supplementary Table 17:** Dunn post hoc test for shoot biomass between control and treated plants

**Supplementary Table 18:** Broad sense heritability values for control (**c**), treated (T) and their ratio (T/Ctrl)

**Supplementary Table 19:** GWAS outcomes

**Supplementary Table 20:** Bacterial and fungal phylum relative abundance across the different compartments

**Supplementary Table 21:** Comparison between the reference sequences used to prepare the inoculum and the sequences detected in the final dataset

**Supplementary Table 22:** Experimental design and leaf phosphate quantification from the additional experiment conducted to validate alpha diversity results

**Supplementary Table 23:** Broad-sense heritability values for ASVs that are differentially abundant in: **a)** plants responding positively vs. negatively to leaf phosphorus (Pi), and **b)** plants with positive vs. negative responses in root biomass

