## Supplementary material for "Plant genetic and root-associated microbial diversity modulate *Lactuca sativa* responsiveness to a soil inoculum under phosphate deficiency": Table 1

**Table 1. List of SNPs significantly associated with phenotypic and physiological traits.**

| **Phenotypc trait** | **heritability** | **Chromosome position** | **chromosome** | **Allelic frequency** | **P-wald** |
| --- | --- | --- | --- | --- | --- |
| Pi T/Ctrl | 0.323 | 117030769 | 4 | 32.3% | 6.786726e-08 |
| Shoot biomass T/Ctrl | 0.326 | 22753367 | 2 | 8.3% | 2.560994e-08 |
| Shoot biomass T/Ctrl | 0.326 | 172536404 | 5 | 49,2% | 1.434627e-08 |
| Head height T/Ctrl | 0.51 | 22753358 | 5 | 49,6 | 1.075417e-24 |
| Head height T/Ctrl | 0.51 | 36408738 | 5 | 49,6 | 1.075417e-24 |
| Head height T/Ctrl | 0.51 | 36408739 | 5 | 49,6 | 1.075417e-24 |
| Head height T/Ctrl | 0.51 | 21725555 | 9 | 49,6 | 1.075417e-24 |
| Head height T/Ctrl | 0.51 | 295749109 | 5 | 50,8 | 4.762413e-13 |
| Head height T/Ctrl | 0.51 | 66640473 | 1 | 49,2 | 6.187546e-12 |
| Head height T/Ctrl | 0.51 | 209962003 | 2 | 49,2 | 6.187546e-12 |
| Head height T/Ctrl | 0.51 | 209962010 | 2 | 49,2 | 6.187546e-12 |
| Head height T/Ctrl | 0.51 | 209962035 | 2 | 49,2 | 1.731641e-11 |
| Head height T/Ctrl | 0.51 | 22753367 | 5 | 49,2 | 6.667104e-11 |
| Head height T/Ctrl | 0.51 | 85252515 | 3 | 49,2 | 1.651884e-11 |
| NPCI | 0.455 | 74322621 | 3 | 5.5 | 2.438137e-08 |
| NPCI | 0.455 | 115636353 | 8 | 5.1 | 2.722674e-08 |
| NPCI | 0.455 | 20535263 | 9 | 5.1 | 3.841432e-08 |
| NPCI | 0.455 | 20602606 | 9 | 5.5 | 4.549761e-08 |
| PSRI | 0.404 | 49514559 | 8 | 11.2 | 2.792331e-08 |
