## Supplementary Figures for "Plant genetic and root-associated microbial diversity modulate *Lactuca sativa* responsiveness to a soil inoculum under phosphate deficiency"

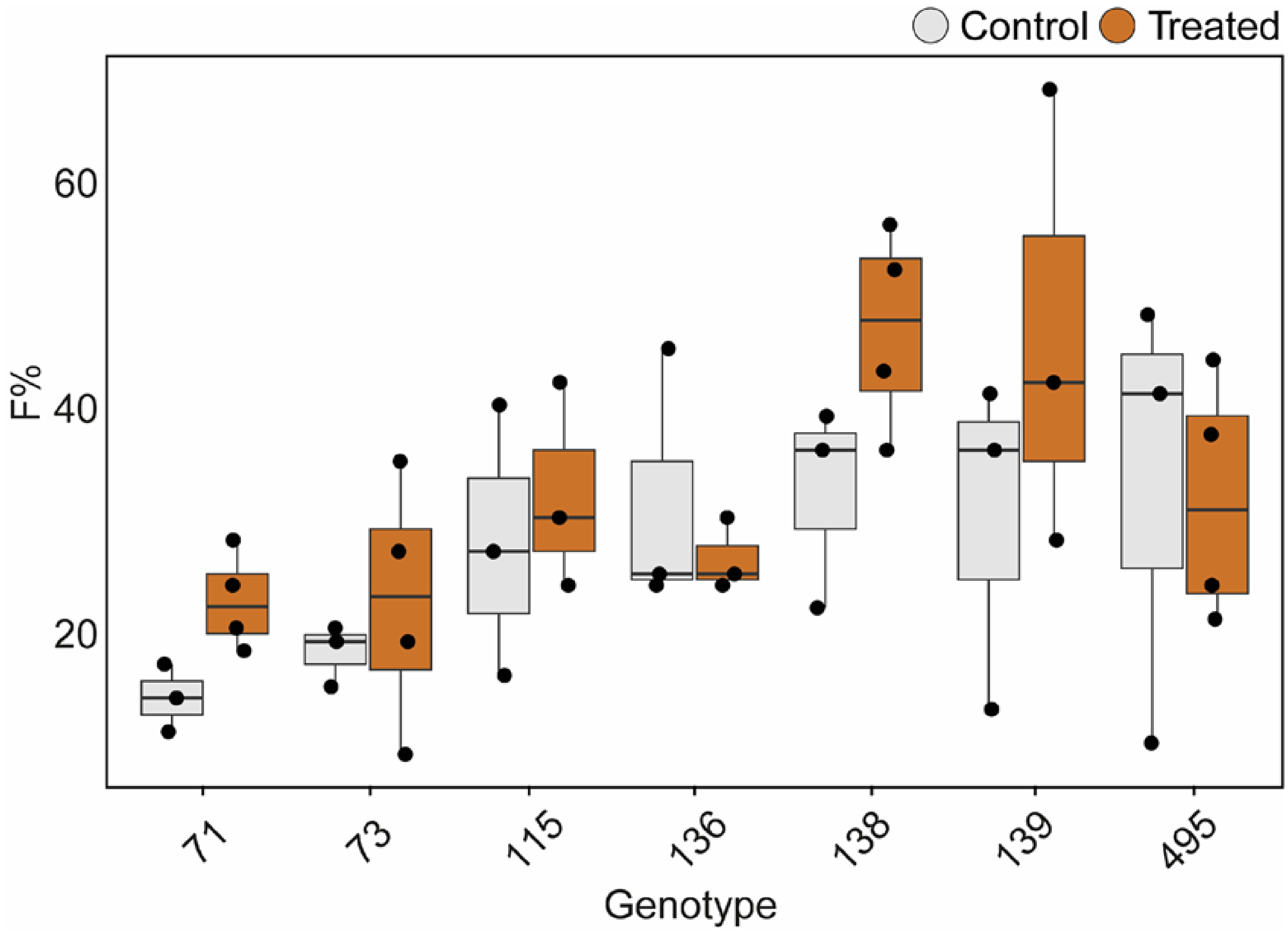


**Fig. S1. Frequency of mycorrhization (F%) in control and treated samples.**

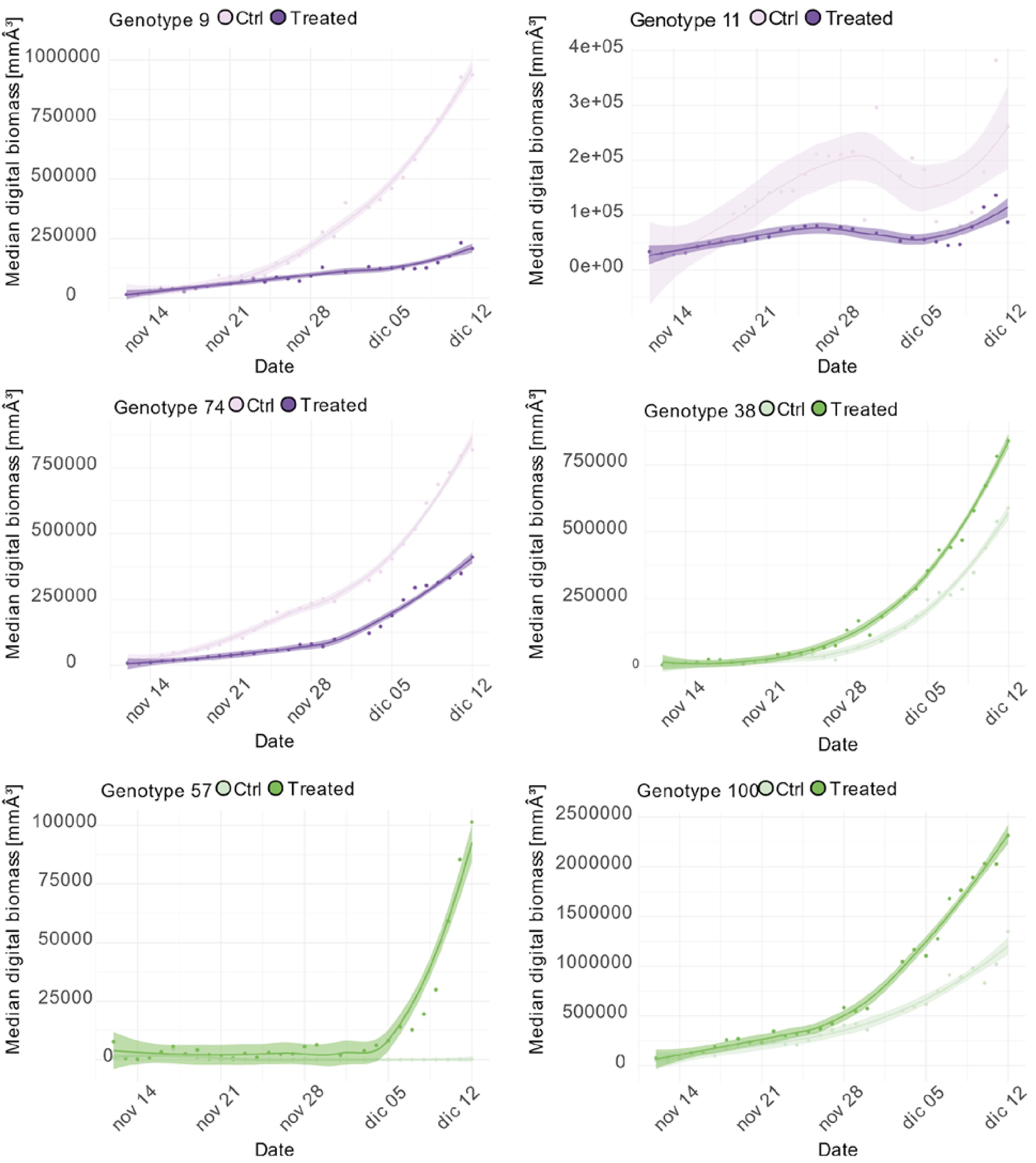


**Fig. S2. Digital biomass trends of genotypes with differential responses (9-11-74-38-57-100) over the course of the experiment.** Genotypes that show a decrease in biomass in response to the treatment are highlighted in purple, while those that experience an increase in biomass due to inoculation are shown in green.

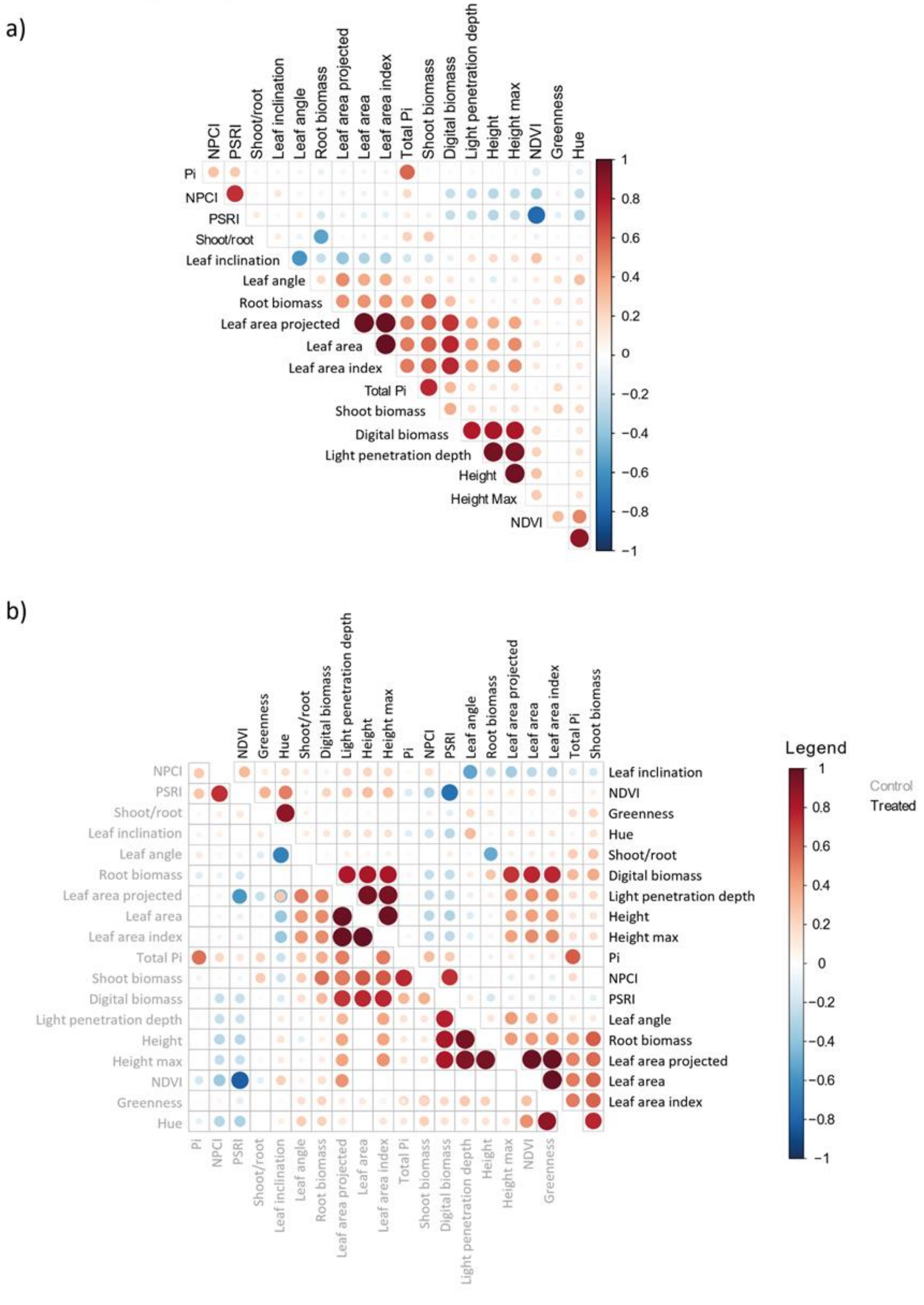


**Fig. S3. Correlation matrix among all parameters measured by the Phenospex system. a)** Correlation matrix including all measured values, and **b)** correlation matrix showing differences between treated and control samples. Positive correlations are colored in red, negative correlations are in blue, and no interactions are in white. Dot size indicates the strength of the correlation, with larger dots representing stronger interactions.

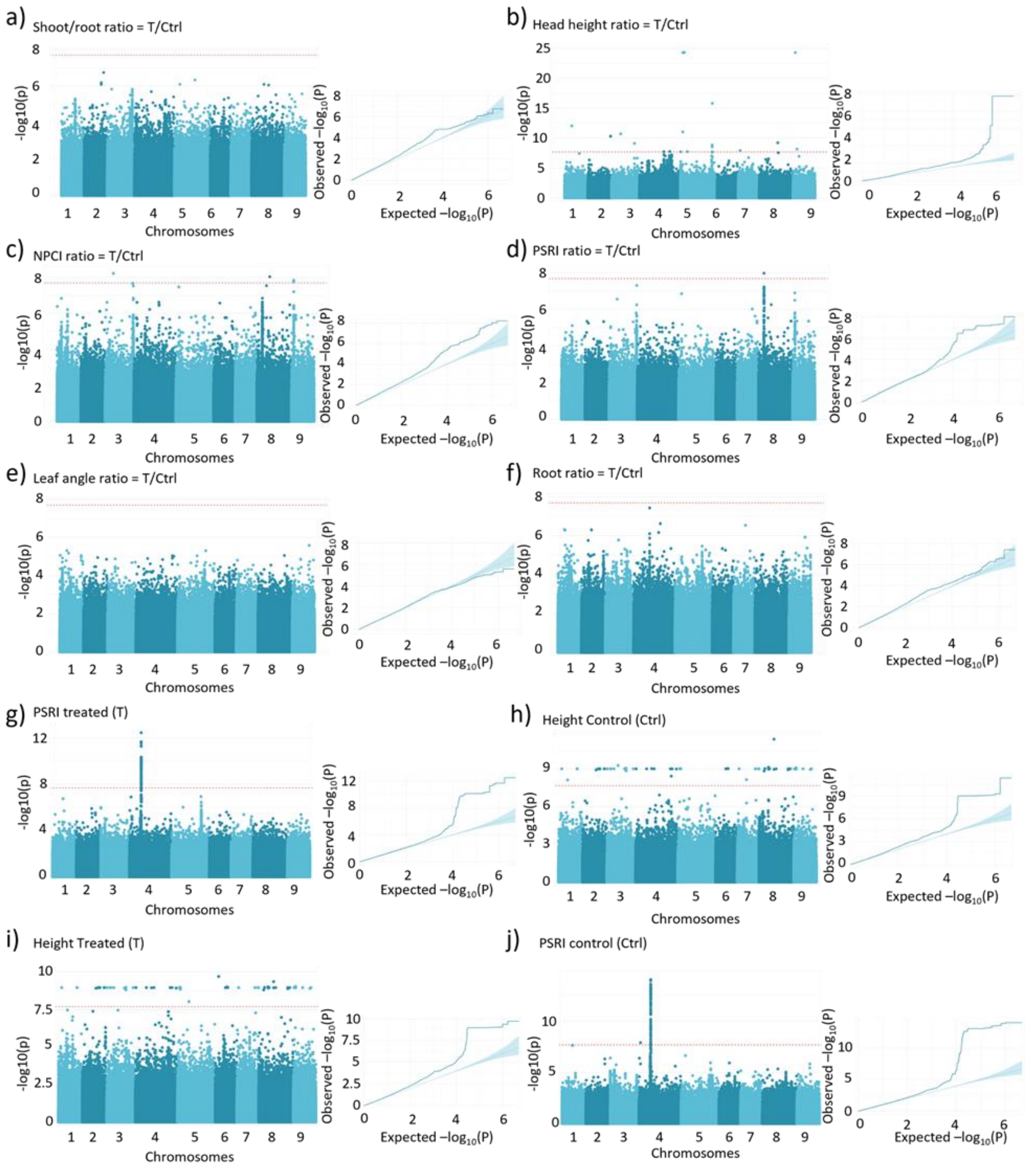


**Fig. S4. GWAS of phenospex parameters**. **a-j**) Manhattan plots and QQ plots showing the correlation between SNPs in the *Lactuca sativa* genome and the following traits: **a)** shoot/root ratio, **b)** head height ratio, **c)** NPCI ratio, **d)** PSRI ratio, **e)** leaf angle ratio, **f)** root ratio, **g)** PSRI treated, **h)** height control, **i)** height treated, **j)** PSRI control

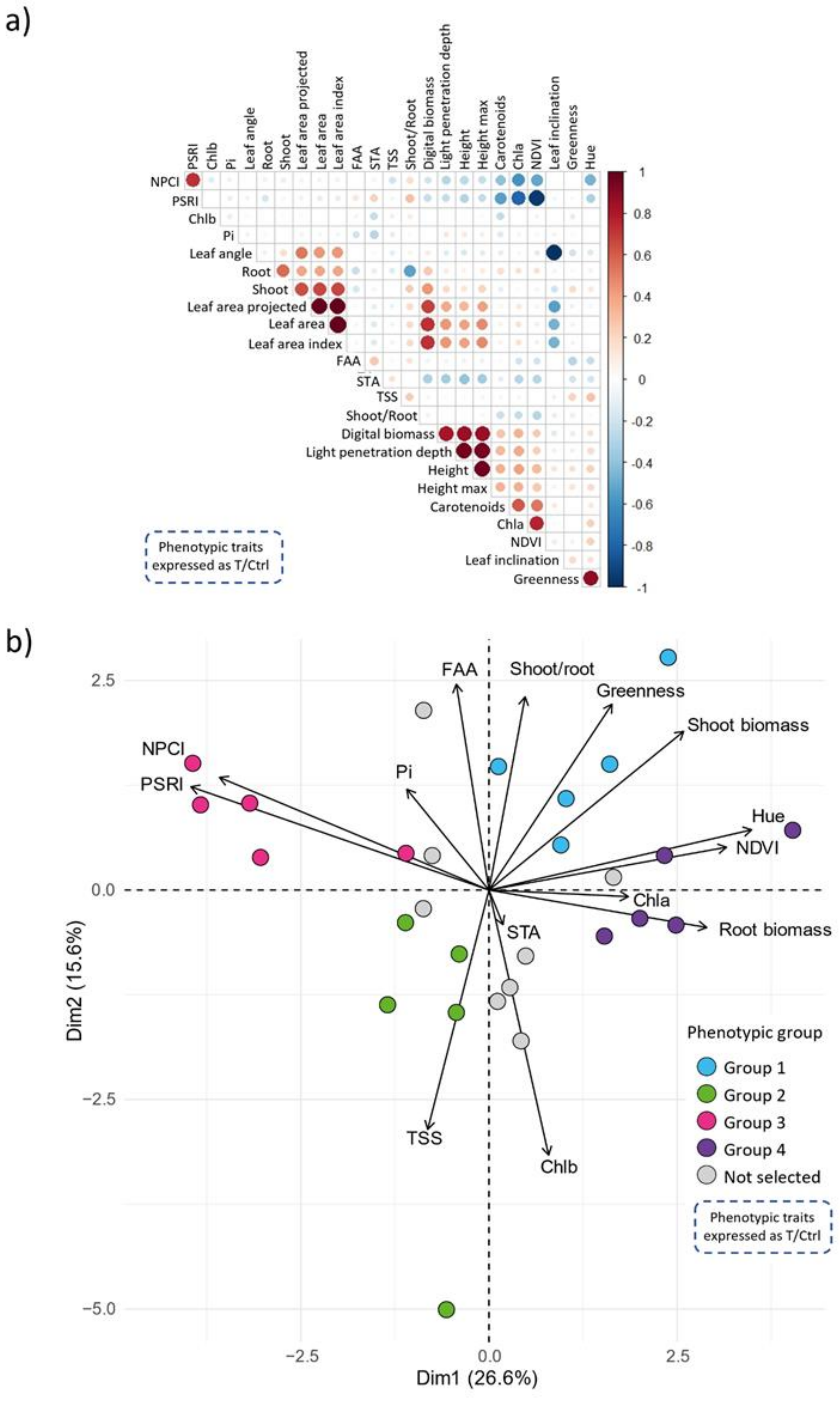


**Fig. S5. Metabolite quantification to validate genotype phenotypic categorization**. **a)** Correlation Matrix among all parameters measured by the Phenospex system and the quantified metabolites. Positive correlations are colored in red, negative correlations are in blue, and no interactions are in white. Dot size indicates the strength of the correlation, with larger dots representing stronger interactions. **b)** Principal component analysis of responses to the inoculum in 28 selected genotypes, based on plant phenotypes and leaf metabolites. Genotypes chosen for root-DNA extraction, representing four distinct phenotypic groups, are color-coded in blue, green, pink, and purple. Non-selected genotypes are displayed in gray.

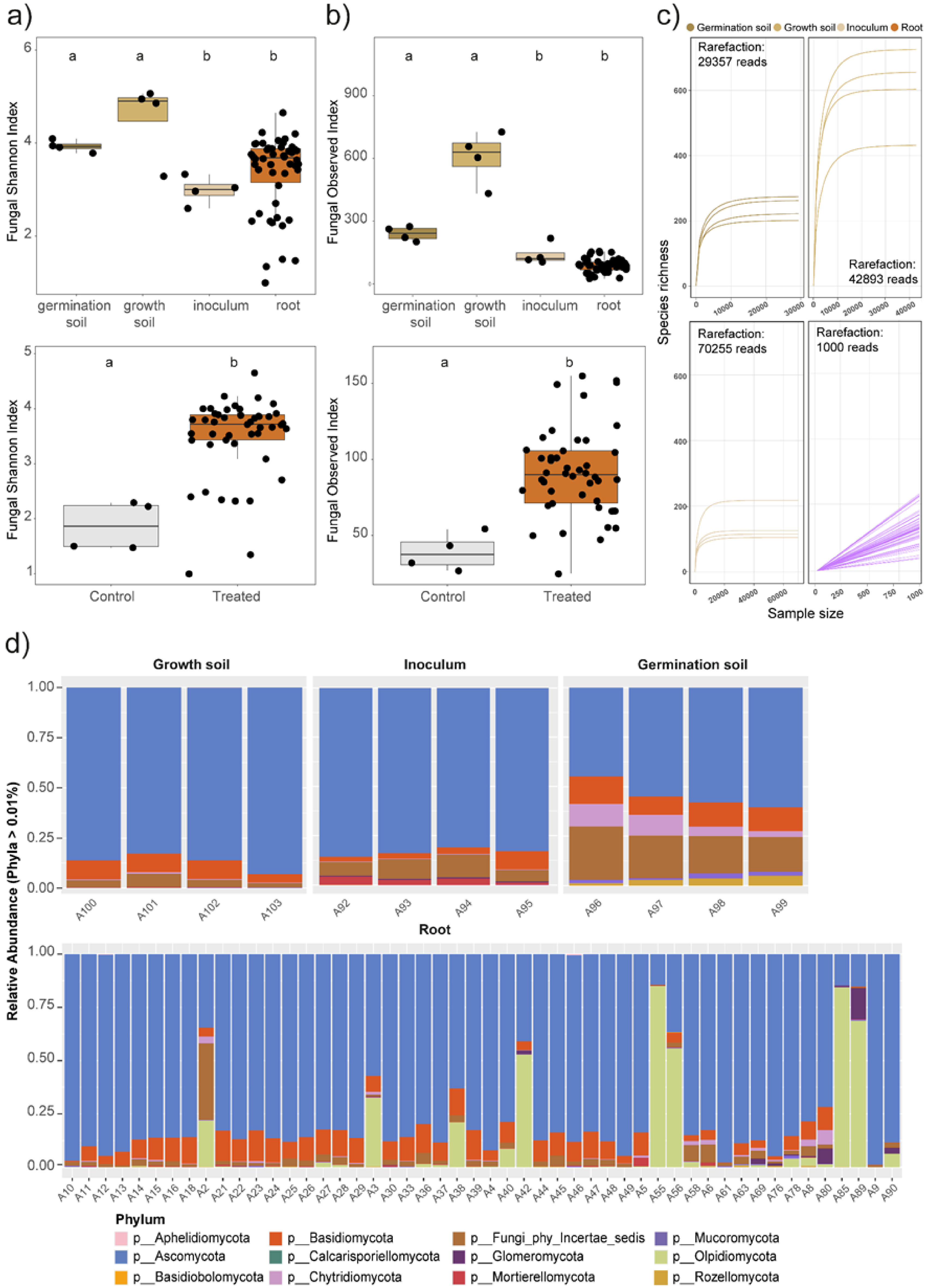


**Fig. S6. Fungal community richness and composition across different compartments.** Panels a,b display Shannon **(a)** and Observed **(b)** α-diversity indices across germination soil, growth soil, inoculum, and root samples. The lower section of each panel compares Shannon **(a)** and Observed **(b)** α-diversity between control (gray) and treated (brown) root samples. **c)** Rarefaction curves following size-based rarefaction across germination soil, growth soil, inoculum, and root samples. **d)** Bacterial community composition at the phylum level across compartments.

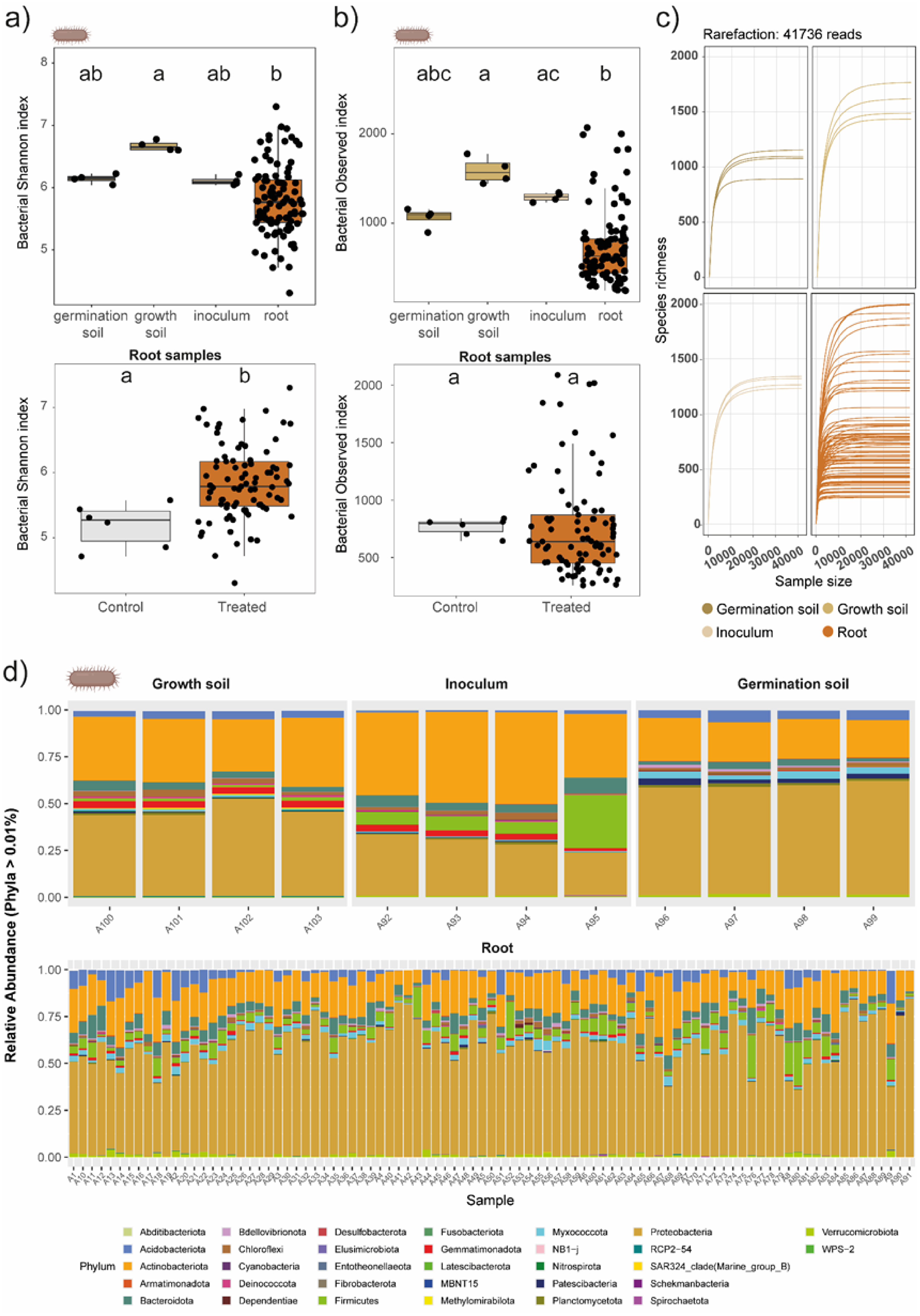


**Fig. S7. Bacterial community richness and composition across different compartments.** Panels a,b display Shannon **(a)** and Observed **(b)** α-diversity indices across germination soil, growth soil, inoculum, and root samples. The lower section of each panel compares Shannon **(a)** and Observed **(b)** α-diversity between control (gray) and treated (brown) root samples. **c)** Rarefaction curves following size-based rarefaction across germination soil, growth soil, inoculum, and root samples. **d)** Bacterial community composition at the phylum level across compartments.

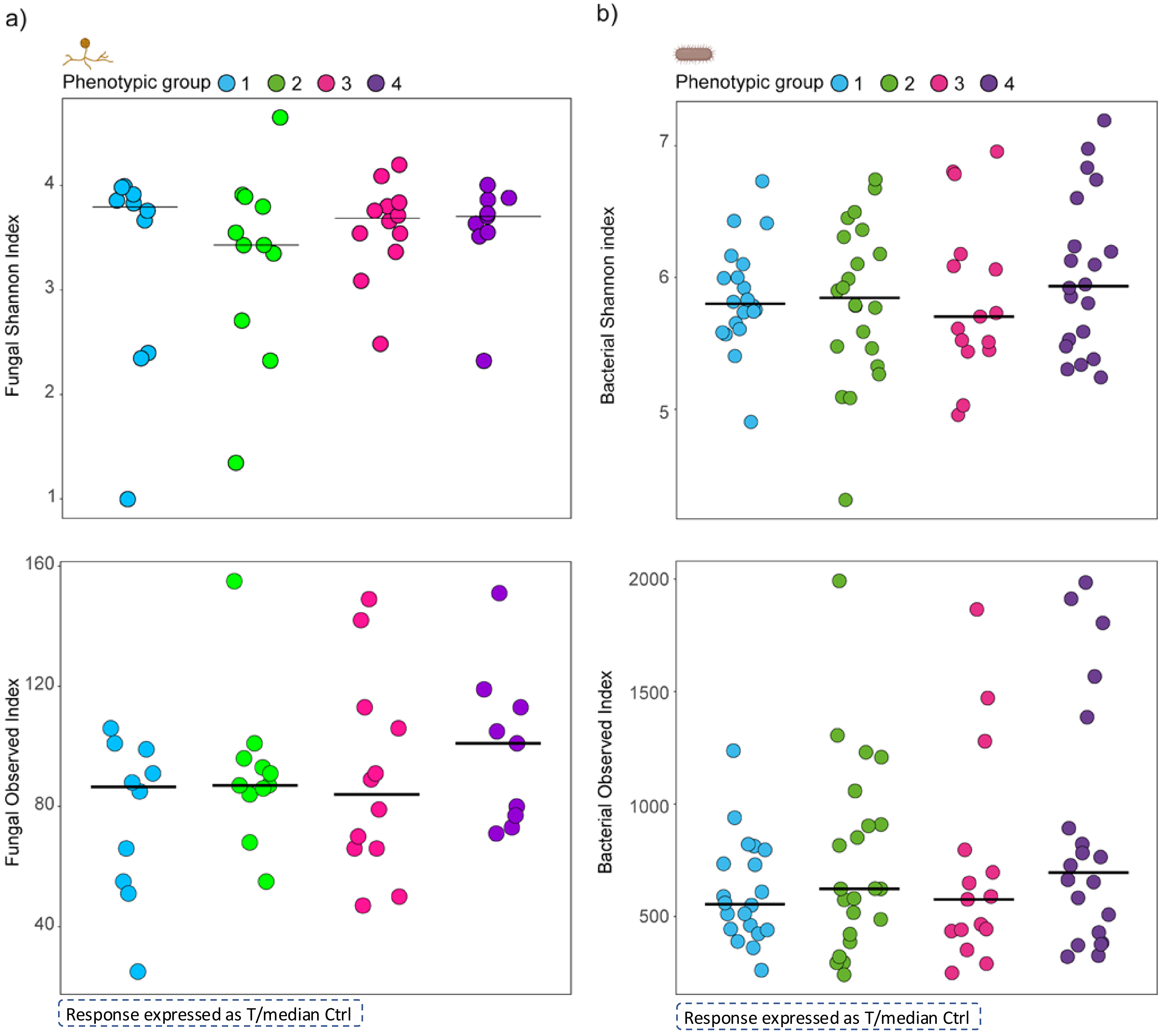


**Fig. S8. Root-associated microbial variation at the base of phenotypic plant response. a)** Fungal and **b)** Bacterial Shannon (above) and Observed (below) α- diversity indices across the four phenotypic groups, indicated by different colors.

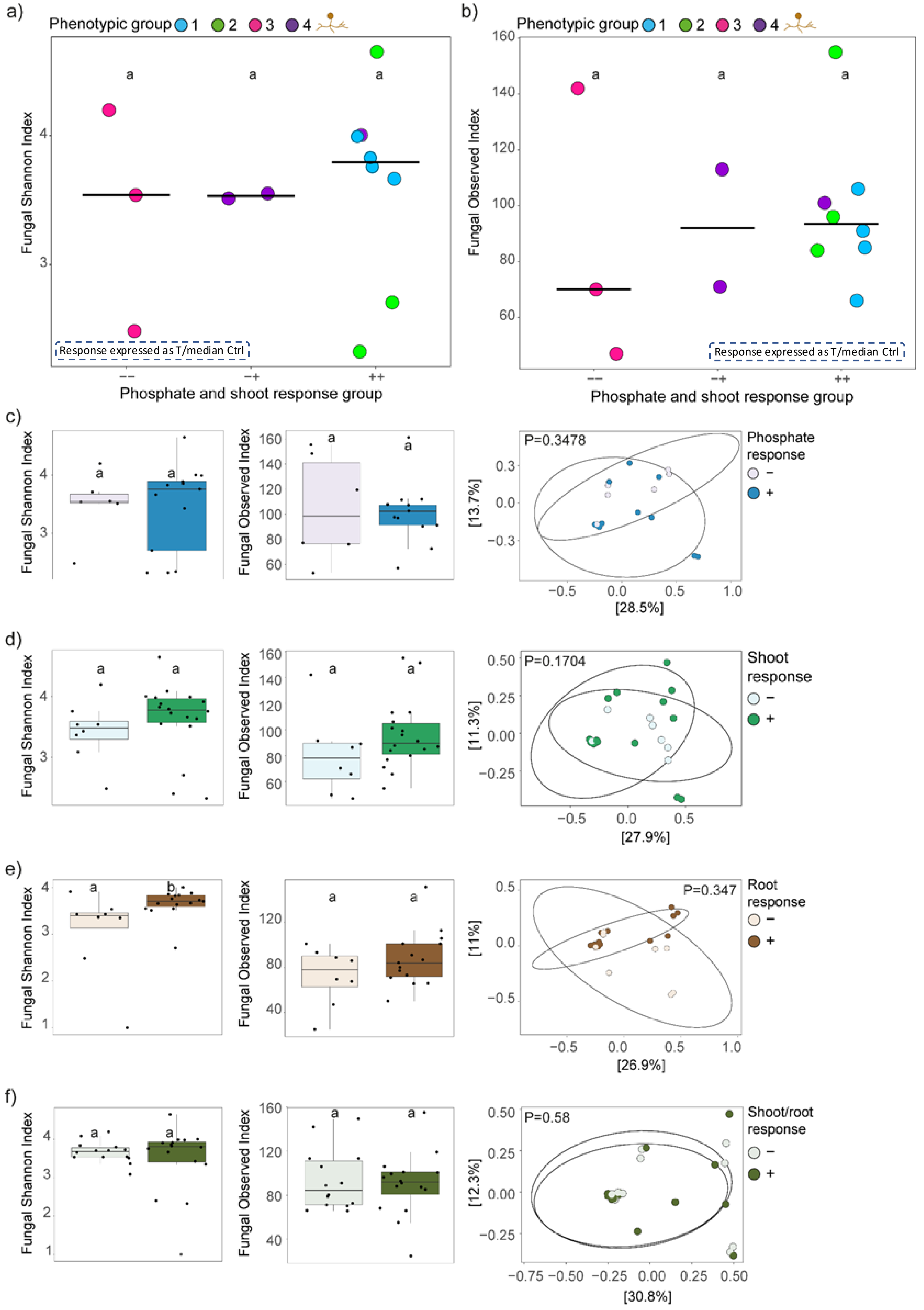


**Fig. S9. Root-associated Fungal community variation at the base of divergent responses**. **a)** Fungal Shannon and **b)** Observed α- diversity indices across the three response groups (--,-+-,++) for leaf phosphate and shoot biomass variation, with each replicate color-coded by the phenotypic group. **c-f)** Fungal α-diversity (Shannon, and Observed indexes) and β- diversity indices comparing positive and negative responses in leaf phosphate accumulation **(c)**, shoot biomass **(d)**, root biomass **(e)**, and shoot/root biomass variation **(f)**.

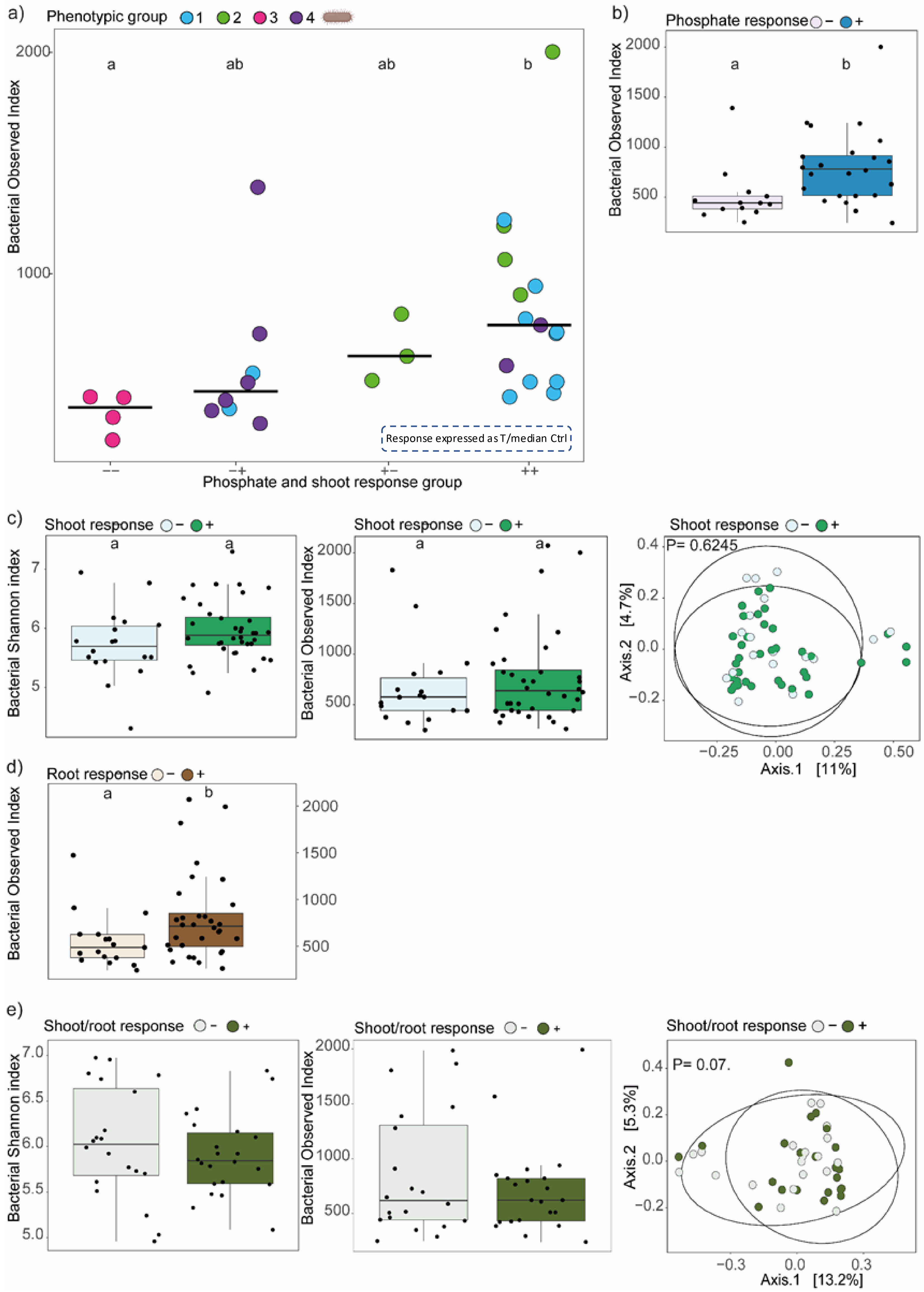


**Fig. S10. Root-associated bacterial community variation at the base of divergent responses**. **a)** Bacterial Observed α-diversity indices across the four response groups (--,-+,+-,++) for leaf phosphate and shoot biomass variation, with each replicate color-coded by the phenotypic group. **b)** Shannon diversity index comparing plants with positive (blue) and negative (gray) responses to leaf phosphate accumulation. **c-f)** Bacterial β-diversity, Shannon, and Observed α-diversity indices comparing positive and negative responses in shoot biomass **(c)**, root biomass **(d)**, and shoot/root biomass variation **(e)**.

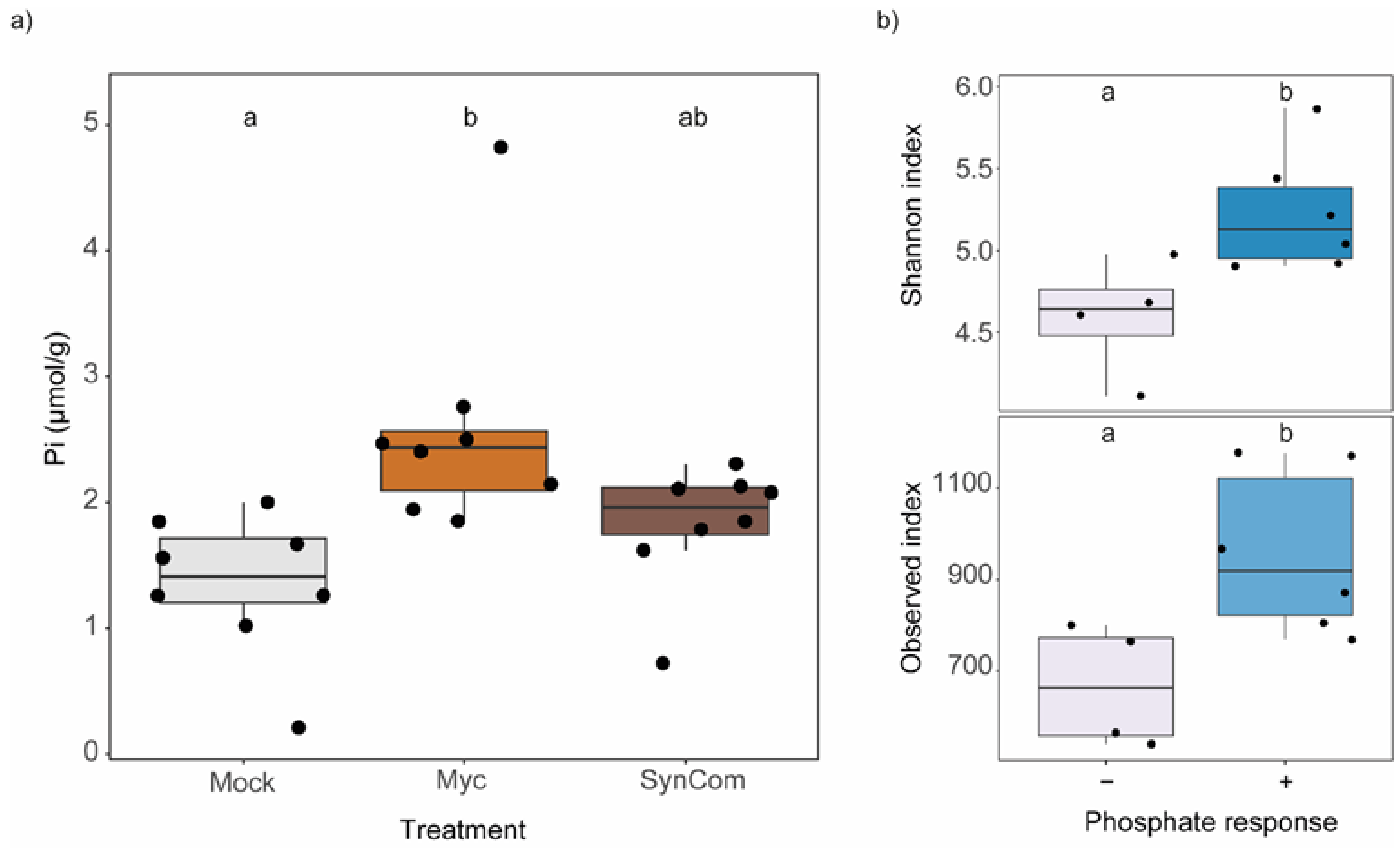


**Fig. S11. Leaf Pi response to fungal (Myc) and synthetic microbial community (SynCom) treatments under Pi deficiency. a)** Difference in soluble phosphate levels between mock-treated plants (gray), plants treated with fungus alone (Myc, light brown), and plants treated with the microbial inoculum (SynCom, dark brown). **b)** Shannon (top) and Observed (bottom) bacterial alpha diversity indices comparing plants with strong (blue) or low (gray) leaf phosphate accumulation compared to mock-treated plants.

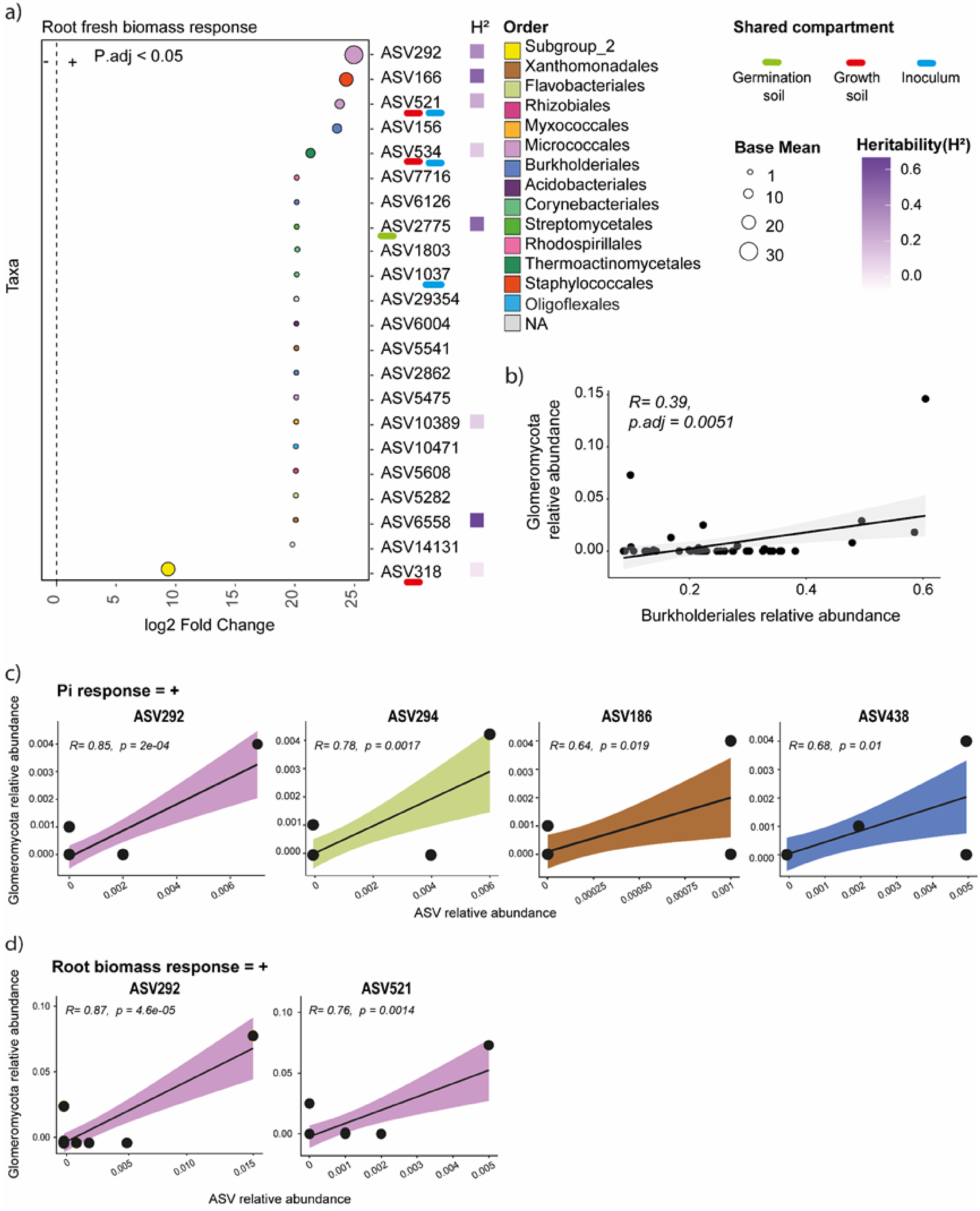


**Fig. S12. Correlation analysis between the root-associated fungal community and the bacterial ASVs enriched in the plants with a positive leaf phosphate and root biomass response. a)** Log2 fold-change values for ASVs, including those with a base mean below 10, showing their abundance in plants with positive (+) and negative (-) variations in root fresh biomass in response to the inoculum. Each dot represents an ASV, with colors indicating its taxonomic order and dot size representing the base mean effect. ASVs shared with germination soil are highlighted in green, those shared with growth soil in red, and those shared with the inoculum in blue. Broad-sense heritability ($H^2$) is shown using a purple color scale, with higher $H^2$ values in dark purple and lower values in lighter shades. **b)** Pearson's correlation between the relative abundance of Burkholderiales (on the x-axis) and the relative abundance of Glomeromycota (on the y-axis). **c-d)** Pearson's correlation between the relative abundance of ASVs 292, 294, 186, 438, and 521 (x-axis) and the relative abundance of Glomeromycota (y-axis). The color of the confidence interval indicates the taxonomic order of each ASV. Panel **c** shows ASVs enriched in plants with a positive phosphate uptake response, while panel **d** shows those enriched in plants with increased root biomass. Only statistically significant correlations are shown.
