## Supplementary materials and methods for "Plant genetic and root-associated microbial diversity modulate *Lactuca sativa* responsiveness to a soil inoculum under phosphate deficiency"

To validate the findings from the main experimental setup, we conducted a smaller-scale experiment using two genotypes, TKI-24 and TKI-57. These genotypes were selected based on their contrasting responses in terms of biomass production and leaf phosphate accumulation observed in the main experiment.

### Experimental design and sample collection

Plant germination, growth, and sampling followed the same conditions and timing as the previous experiment but were conducted in a growth chamber at the Department of Biology, University of Padova. For the treatment, in addition to using the same microbial consortium described in the main experiment, we included a second commercial inoculum (Agronutrition, Labege, France) containing only arbuscular mycorrhizal fungus propagules (*Rhizophagus irregularis* - DAOM 197198) Specifically, one-month-old seedlings were transplanted into a soil–sand mixture (5 L of LS mix sterilized in a dry stove at 190 °C for 1 h) supplemented with pre-weighed triple superphosphate (TSP) at the same concentration used in the previous setup, designed to create phosphorus-deficient conditions. Plants were arranged in randomized positions and treated immediately after transplantation. Due to germination failure, only 9 seedlings were available for genotype TKI-57 and 15 for genotype TKI-24. These included mock plants (5 replicates for TKI-24, 3 for TKI-57), plants treated with the fungal inoculum (5 for TKI-24, 3 for TKI-57), and plants treated with the synthetic microbial community (5 for TKI-24, 3 for TKI-57).

### Leaf Phosphate quantification and DNA extraction from roots

After approximately one month of growth, leaf and root samples were collected. For leaf analysis, three leaf disks per plant were pooled, dried at 70°C overnight, and homogenized for soluble phosphate quantification. Fresh root samples were first thoroughly washed to remove residual sand and subsequently processed for DNA extraction using the DNeasy® PowerSoil® Kit (QIAGEN, Hilden, Germany), following the manufacturer's instructions. The extracted DNA was then used to characterize the root-associated bacterial community via 16S rRNA sequencing. Procedures for phosphate quantification, DNA sequencing, and bacterial community analysis followed the protocols described in the main Materials and Methods section.

The experimental setup and phosphate quantification results are presented in Supplementary Table **19**.

**Statistical analysis**

To analyze the plant responses to treatments in terms of leaf phosphate concentration, each replicate's response was calculated as described in *Equation 8*. This calculation yielded values ranging from 0.69 to 3.13. We then categorized plants based on the overall response range: samples with a weak response (values between 0.68 and 1.25) and those with a strong response (values above 1.6). These groups were then compared for differences in alpha diversity using R Studio. Differences between treated and control plants were assessed using the Kruskal-Wallis test, followed by Dunn's *post hoc* test.
